# A key role of the WEE1-CDK1 axis in mediating TKI-therapy resistance in FLT3-ITD positive acute myeloid leukemia patients

**DOI:** 10.1101/2022.05.16.492070

**Authors:** Giorgia Massacci, Veronica Venafra, Sara Latini, Valeria Bica, Giusj Monia Pugliese, Felix Klingelhuber, Natalie Krahmer, Thomas Fischer, Dimitrios Mougiakakos, Martin Boettcher, Livia Perfetto, Francesca Sacco

**Affiliations:** Department of Biology, University of Rome Tor Vergata, Via della Ricerca Scientifica 1, 00133 Rome, Italy; Institute for Diabetes and Obesity, Helmholtz Zentrum München, 85764 Neuherberg, Germany; German Center for Diabetes Research, Neuherberg, Germany; Institute of Molecular and Clinical Immunology, University of Madgeburg, Madgeburg, Germany; Healthcampus for Inflammation, Immunity and Infection (GCI3), University of Magdeburg, Germany; Department of Hematology and Oncology, University of Magdeburg, Magdeburg, Germany; Fondazione Human Technopole, Department of Biology, Via Rita Levi-Montalcini 1, 20157 Milan, Italy; TIGEM Institute, Via dei Campi Flegrei, Naples, Italy

## Abstract

Internal tandem duplications (ITDs) in the FLT3 gene are frequently identified and confer a poor prognosis in patient affected by acute myeloid leukemia (AML). The insertion site of the ITDs in FLT3 significantly impacts the sensitivity to tyrosine kinase inhibitors (TKIs) therapy, affecting patient’s clinical outcome. To decipher the molecular mechanisms driving the different sensitivity to TKIs therapy of FLT3-ITD mutation, we used high-sensitive mass spectrometry-based (phospho)proteomics and deep sequencing. Here, we present a novel generally-applicable strategy that supports the integration of unbiased large-scale datasets with literature-derived signaling networks. The approach produced FLT3-ITDs specific predictive models and revealed a crucial and conserved role of the WEE1-CDK1 axis in TKIs resistance. Remarkably, we found that pharmacological inhibition of the WEE1 kinase synergizes and strengthens the pro-apoptotic effect of TKIs therapy in cell lines and patient-derived primary blasts. In conclusion, this work proposes a new molecular mechanism of TKIs resistance in AML and suggests a combination therapy as option to improve therapeutic efficacy.

## Introduction

Internal tandem duplications (ITDs) of the FLT3 gene are observed in about 25% of young adults with newly diagnosed acute myeloid leukemia (AML) ^1,2^. The FLT3 gene encodes a receptor tyrosine kinase, consisting of an extracellular immunolike-domain, a transmembrane region, a cytoplasmic juxtamembrane domain (JMD) followed by two tyrosine kinase domains (TKD1 and TKD2) ^3^. Upon ligand binding, FLT3 switches from an inactive to a catalytically active conformation, leading to the phosphorylation of its downstream effectors that regulate self-renewal and differentiation of hematopoietic stem and progenitor cells. FLT3-ITD mutations always occur in exons 15 and 16, encoding the JMD and TKD1 regions, and cause its constitutive activation ^4^. In 2017, the CALGB 10603/RATIFY trial demonstrated a significantly improved outcome in a cohort of 717 patients carrying genetic alterations in the FLT3 gene when treated with the multikinase inhibitor midostaurin (PKC412) combined to standard frontline chemotherapy ^5^. At the beginning of 2022, a retrospective analysis of the same trial evaluated the prognostic value of the insertion sites of ITD mutations in the response to midostaurin treatment. Interestingly, the analysis revealed that midostaurin treatment exerted a significant beneficial effect only in patients carrying the ITDs in the JMD domain, whereas no beneficial effect was observed in patients carrying ITDs in the TKD region. In addition, multivariate analysis showed that the ITD-TKD localization is an unfavourable prognostic factor for overall survival and incidence of relapse ^6^.

In accord with this clinical observations, previous *in vitro* studies showed that ITDs-TKD confer resistance to chemotherapy and are associated to a significantly inferior outcome. Briefly, ITD-TKD positive cell lines and primary mouse bone marrow cells showed reduced apoptosis when compared to ITDs-JMD, upon exposure to FLT3 inhibitors, namely midostaurin and quizartinib (a highly-specific second-generation FLT3 inhibitor) ^7–9^.

Although ITD-TKD and ITD-JMD expressing cell models show two clearly distinct phenotypes, at the biochemical level, the enzymatic activity of these FLT3 mutants is equally suppressed by kinase inhibitors. In addition, downstream FLT3 canonical targets are also equally inhibited upon drug administration ^7^, suggesting that the two differential phenotypic outcomes are the result of a systems-wide response to inhibitor treatment.

To date, the molecular mechanisms underlying such different sensitivity remain unclear, leaving patients carrying ITDs in the TKD region in still unmet medical needs.

Here, we speculate that the diverse insertion site of ITDs in FLT3 may cause an extensive rewiring of cell signaling network resulting in a different sensitivity to TKIs therapy. A variety of systems biology and network-based approaches have been recently developed to generate predictive cell specific models ^10^. In principle cell specific signaling models describing how the different ITD localization impacts the signaling network may offer the opportunity to identify novel promising therapeutic targets reverting the TKI-therapy resistance ^11,12^. We, therefore, performed a system-level analysis of the state of FLT3-ITD cells caused by TKIs treatment. Global and unbiased transcriptome, proteome and phosphoproteome analyses enabled us to identify TKIs-induced changes at transcriptional, translational and post-translational levels. Next, to obtain FLT3-ITD cell-specific models describing the TKI signaling response, we developed “Signaling Profiler”, a novel, generally applicable, computational strategy supporting the integration of these large “omic” datasets with literature-derived causal networks. This strategy highlighted the novel and crucial role of the WEE1-CDK1 axis in TKI therapy failure in FLT3^ITD-TKD^ patients. Remarkably, pharmacological inhibition of WEE1 completely rescued the ability of patient-derived primary blasts, carrying the ITD-TKD mutation, to undergo apoptosis in response to midostaurin treatment. Thus, our strategy and resource enable investigation of drug-resistance mechanisms and identification of therapeutic targets.

## Results

### An unbiased approach to identify therapeutic target reverting TKI resistance

In our experimental strategy we aim to dissect the molecular mechanisms underlying the different impact of the location of ITDs on sensitivity to FLT3 inhibitor treatment, with the ultimate goal of designing effective patient-specific therapeutic strategies.

To this aim, we set out a multi-step strategy that combines system-level and unbiased multi-omic analyses (**Fig.1A** **panel a and b**) with literature-derived causal networks to generate cell-specific models (**Fig.1A** **panel c**). We demonstrate that these models have a translational impact and can be used as a framework to identify and test novel drug targets reverting TKI resistance (**Fig.1A** **panel d**).

**Figure 1.**
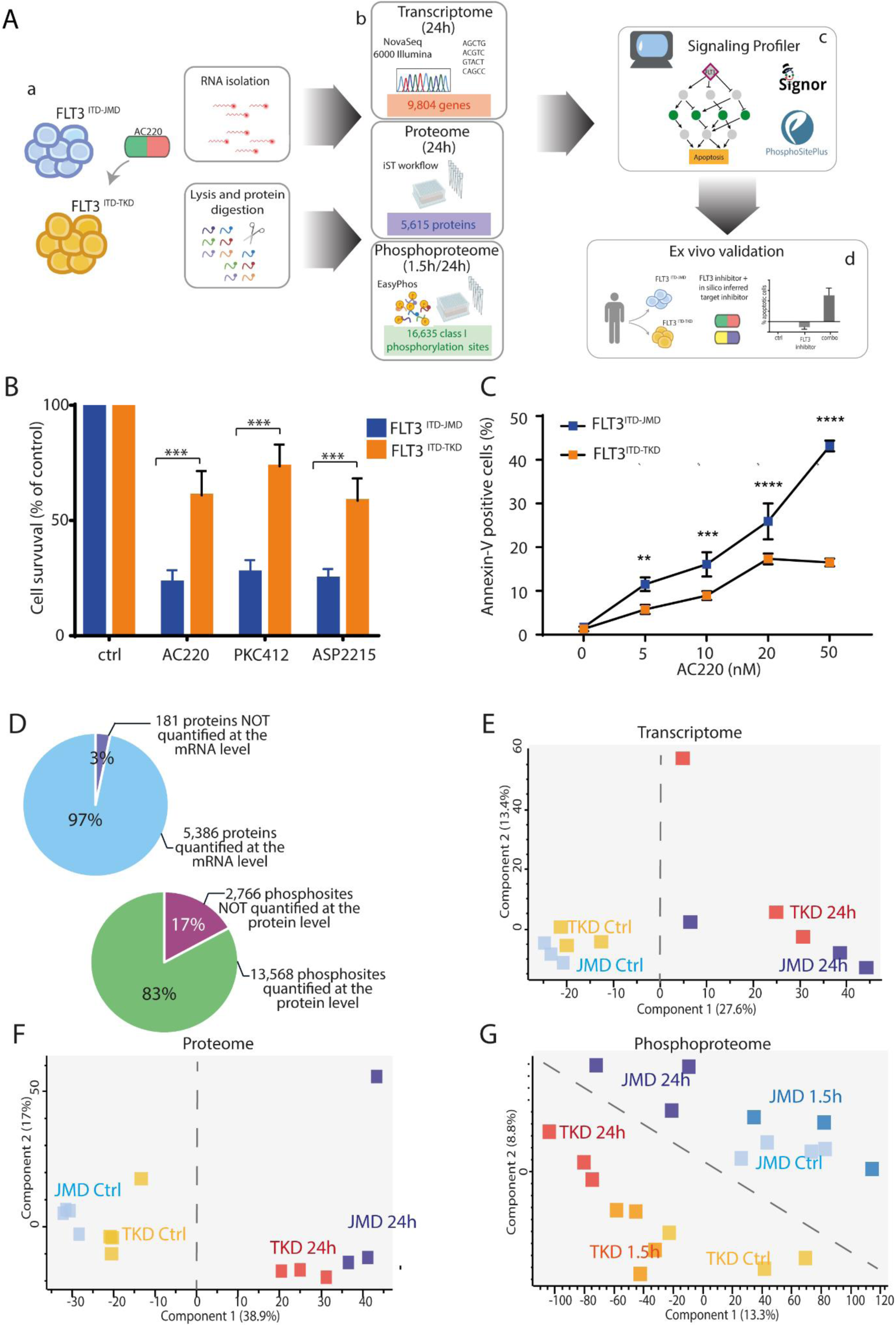
(**A**) Overview of the experimental and bioinformatic strategy. BaF3 cells expressing FLT3^ITD-TKD^ (in orange) and FLT3^ITD-JMD^ (in blue) were treated with 20nM quizartinib (AC220) for 24h (a). mRNAs were isolated for the transcriptome analysis and the protein extracts were digested and characterized at the proteome and phosphoproteome levels (b). Multi-omics profiles of FLT3-ITD cells were used in *Signaling Profiler* pipeline to obtain cell-specific models and to identify additional druggable genes (c). Proteins of interest were further investigated through complementary assays in patients-derived primary blasts (d). (**B**) Cell survival of BaF3 cells expressing FLT3^ITD-JMD^ (in blue) and FLT3^ITD-TKD^ (in orange) after FLT3 inhibitors treatment. Cells were treated for 24h with 20nM quizartinib (AC220), 100nM midostaurin (PKC412) and 50nM gilteritinib (ASP2215). Cell viability was assessed by MTT assay. (**C**) Induction of apoptosis in BaF3 cells expressing FLT3^ITD-JMD^ (in blue) and FLT3^ITD-TKD^ (in orange) treated with increasing doses of AC220 for 24h. The percentage of apoptotic cells was determined by Annexin-V labelling. (**D**) Pie charts representing the percentage and the number of species characterized at the protein and the transcript levels (top) or at the protein and the phosphosite levels (bottom). (**E, F, G**) Principal Component Analysis (PCA) of the analytes quantified across the transcriptome (**E**), proteome (**F**) and phosphoproteome (**G**) replicates.

### The experimental system

Specifically, as an experimentally easy-to-manipulate system, we used Ba/F3 cells stably expressing the FLT3 gene with ITD insertions in the JMD (aa 598) or in the TKD1 (aa 613) region, from now on “FLT3^ITD-JMD^” and “FLT3^ITD-TKD^” cells (**Fig. 1A** **panel a**). Among validated FLT3 inhibitors, we tested midostaurin (PKC412), gilteritinib (ASP122, recently approved by FDA) and quizartinib (AC220) ^13 14^.

As anticipated, nanomolar concentrations (20–100 nM) of midostaurin (PKC412), quizartinib (AC220) or gilteritinib (ASP122) induced apoptosis in BaF3-cells harboring ITD-JMD mutation, whereas ITD-TKD cells showed significantly decreased sensitivity to the three inhibitors (**Fig. 1B**). A dose-dependent assay confirmed this observation: increasing concentrations of quizartinib, the more selective and potent FLT3-inhibitor ^15^, correlate with a stronger response in ITD-JMD cells (**Fig. 1C**). Confirming previously published data ^7,8^, these results demonstrate differential sensitivity of FLT3-ITDs to TKI-therapy depending on the location of the FLT3-ITD.

### Deep transcriptome, proteome and phosphoproteome analysis of quizartinib treated FLT3-ITD cells

To investigate the molecular basis of the observed different sensitivities to treatment with FLT3 inhibitors in ITD-TKD and ITD-JMD expressing cells, we set out to apply an unbiased strategy to monitor the transcriptional, translational, and post-translational changes induced by FLT3 inhibition. In these large-scale experiments, FLT3-ITD cells were exposed to short (1.5h) or long (24h)-term treatments with 20nM quizartinib (AC220) (**Fig. 1A** **panel b**). Among the FLT3 inhibitors, we selected quizartinib, because of its high specificity and we treated cells with 20nM, a non-toxic concentration whereby the quizartinib-induced apoptotic response is significantly different between FLT3-ITD cells (**Fig. 1C**).

We quantified the modulation of the transcription profiles induced by TKI in FLT3-ITD cells. A deep sequencing approach enabled the quantification of the expression of more than 11,000 genes (**Table S1**). Protein levels and peptide phosphorylation were obtained by state-of-the-art, high-sensitive, mass spectrometry (MS)-based (phospho)proteomics. This allowed the quantification of more than 5,000 proteins (**Table S2)** and 16,000 phosphorylation events (class I sites, **Table S3**). Overall, by this strategy we quantified more than 10,000 transcripts, 4,000 proteins and 10,000 phosphosites in each experimental condition (**Fig. S1 A-B-C**). The biological triplicates or quadruplicates were highly correlated with Pearson correlation coefficients ranging between 0.85 (for phospho measurements) and 0.97 (for transcriptome and proteome measurements) (**Fig. S1 D-E-F**).

The experimental system appeared to be efficient: for nearly all the quantified proteins (97%), we also obtained the levels of the corresponding transcripts (**Fig. 1D** **top panel**); similarly, for approximately 83% of the quantified phosphorylation sites, we also measured protein abundance (**Fig. 1D** **bottom panel**); also, quizartinib-induced changes at the transcript level tend to correlate with those at the proteome level in FLT3^ITD-JMD^ and FLT3^ITD-TKD^ cells (PC=0.6-0.7) (**Fig. S2 A-B**); finally, when normalizing by the protein levels, more than 70% of phosphosites were still significantly regulated by quizartinib treatment (**Fig. S2 C-D**).

Next, we applied a statistical t-test to narrow-down the species that are regulated by the FLT3 inhibitor. Briefly, about one third of the transcriptome, proteome and phosphoproteome displayed a significant (FDR<0.1) change in the abundance upon quizartinib treatment (**Fig. S3 A-B-C**). Comparative analysis of significantly modulated genes, proteins or phosphosites, revealed a common core (14% in the transcriptomics, 7% in the proteomics and 5% in the phosphoproteomics) of canonical FLT3-ITD targets significantly altered by 24h quizartinib treatment, regardless of the ITD insertion site (**Fig. S3 A-B-C**). Interestingly, our data suggest that the two different ITD localization impacts the quizartinib-dependent remodeling of the phosphoproteome profile to a greater extent as compared to the transcriptome and proteome profile (R phosphoproteome= 0.58) (**Fig. S3 D-F**).

Consistently, principal component analysis (PCA) clearly showed that exclusively the phosphoproteome could stratify cells according to both FLT3 activation status (component 1) and ITD insertion site (component 2) (**Fig. 1 E-G**). Unsupervised hierarchical clustering of our large-scale datasets confirmed that the phosphoproteomic profile best discriminates FLT3 cells according to their quizartinib sensitivity (**Fig. S3 G-I**). These observations indicate that the different localization of the ITD mutations mostly impact the cell regulatory network at the post-translational level. The FLT3-ITD-dependent modulation of the phosphoproteome profile may drive the different sensitivity to TKI-therapy.

### Pathway modulation in response to quizartinib treatment

We next assessed the effect of quizartinib treatment on previously identified FLT3 downstream signaling pathways. We took advantage of the signaling database SIGNOR ^16^ to retrieve the subnetwork recapitulating well-characterized phosphorylation events directly or indirectly modulated by the FLT3 receptor. We, next, overlaid our phosphoproteomic results onto the FLT3 subnetwork. As shown in **Figure 2A**, the MAPK and AKT-mTOR pathways are equally inhibited by either short-term and long-term exposure to quizartinib in both cell lines. These results are in line with literature-derived prior knowledge and support the robustness of our experimental model. Consistently, the phosphorylation level of the FLT3-ITD mutants as well as their down-stream canonical targets are equally decreased by TKIs treatments (**Fig. 2B**).

**Figure 2.**
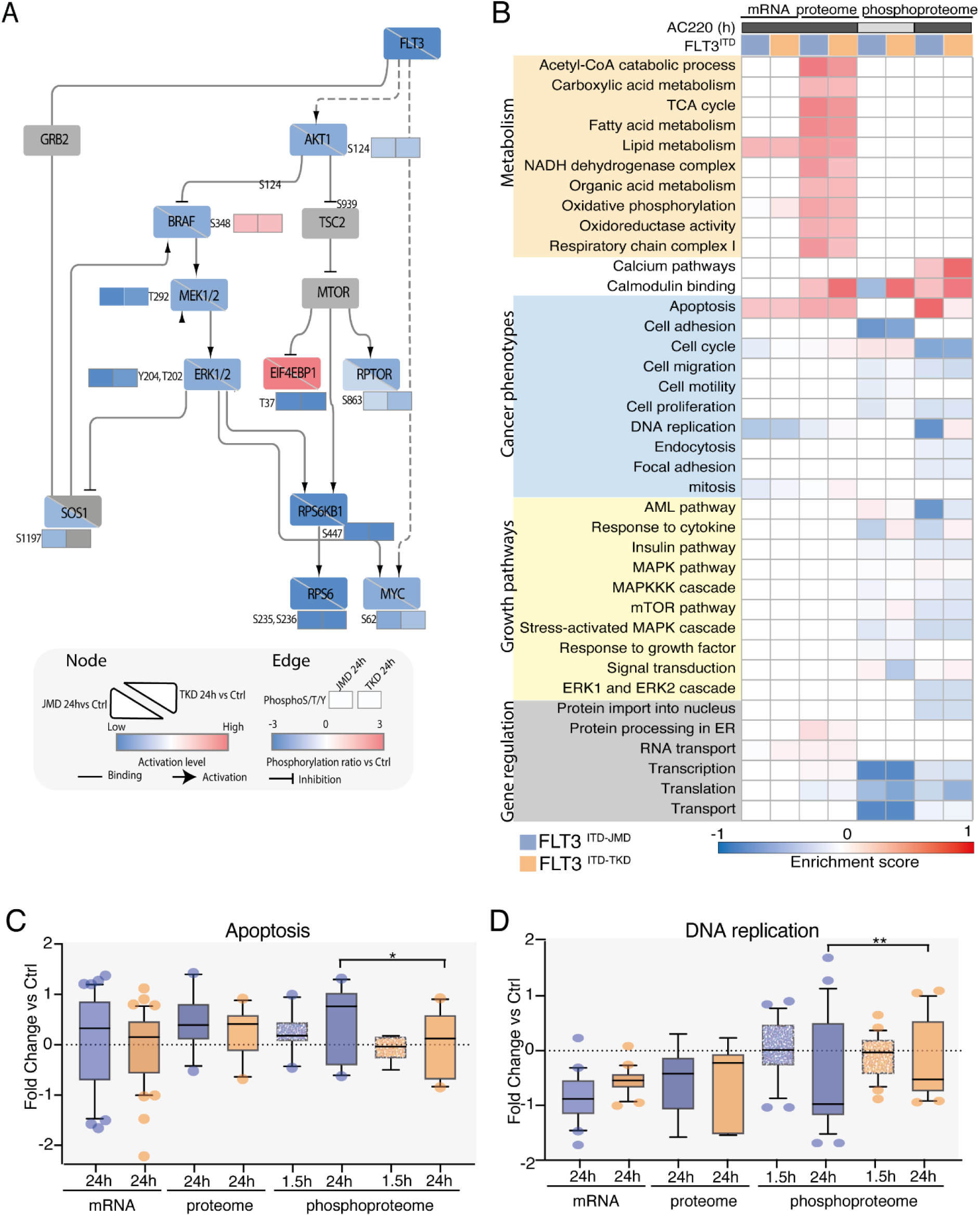
(**A**) FLT3 downstream causal interaction network. The effect of quizartinib (AC220) on the phosphoproteome and proteome profiles of FLT3^ITD-JMD^ and FLT3^ITD-TKD^ cells was mapped on a literature curated signaling network, extracted from the SIGNOR resource ^16^. For comparative analysis, for each node, the activation state in both cell line is shown (down-left half for ITD-JMD, and top-right half for ITD-TKD). Activated proteins are marked in red, whereas inhibited ones in blue. Phosphosites are displayed as independent rectangles and are colored according to their phosphorylation state after quizartinib treatment, as indicated in the legend. (**B**) Representative western blot showing the inhibition of canonical FLT3 downstream targets, as revealed by their phosphorylation status: Tyr694 in STAT5 and Thr202 and Tyr204 in ERK1/2. ITD-JMD and ITD-TKD BaF3 cells were treated for 1.5h with FLT3 inhibitors treatment (AC220: quizartinib, PKC412: midostaurin and ASP2215: gilteritinib). (**C**) Heatmap displaying the enrichment score of GO Biological processes and KEGG pathways significantly (FDR<0.05) over- (red color) or under- (blue color) represented in the relative dataset (transcripts, proteins and phosphosites in FLT3^ITD-JMD^ (in blue) and FLT3^ITD-TKD^ (in orange) BaF3 cells upon quizartinib (AC220) treatment. (**D**, **E**) Boxplots showing the relative abundance of significantly modulated transcripts, proteins and phosphoproteins involved in apoptosis (**D**) or DNA replication process (**E**) in BaF3 cells expressing FLT3^ITD-JMD^ (in blue) and FLT3^ITD-TKD^ (in orange) upon quizartinib (AC220) treatment.

One step further, we checked for biological processes altered in treated cells by employing a gene ontology term enrichment analysis and by taking advantage of the 2D annotation enrichment analysis algorithm, based on nonparametric Mann-Whitney test (FDR < 0.05). We detected a significant overexpression of proteins involved in mitochondrial metabolic processes, such as TCA cycle and OxPhos, as well as in lipid oxidation (**Fig. 2C** **and Fig. S4A-E**).

Consistently with the different sensitivity of FLT3^ITD-JMD^ and FLT3^ITD-TKD^ cells to quizartinib treatment we found that proteins involved in apoptosis were significantly hyperphosphorylated in FLT3^ITD-JMD^ cells, but not in FLT3^ITD-TKD^ cells (**Fig. 2D**). Interestingly, we observed that the quizartinib-dependent phosphorylation of DNA replication proteins is significantly decreased only in quizartinib treated FLT3^ITD-JMD^ cells, but not in cells with the ITD mutation in the TKD region (**Fig. 2E**).

Kinase substrate motifs analysis showed that pro-proliferative kinases, ERK1/2, AKT and p70S6K are significantly downregulated (FDR < 0.05), in line with the anti-proliferative effect of quizartinib in FLT3-ITD cells (**Fig. S4 F**).

These observations provide a global picture of the main changes induced by quizartinib treatment at the transcriptome, proteome, and phosphoproteome level in both FLT3-ITD cells. The molecular mechanisms underlying the different sensitivity to quizartinib treatment of FLT3-ITD cells is still an open question that we aim to address by complementary and more granular approaches.

### From FLT3 to transcription factors through *Signaling Profiler*

Here we implemented a generally applicable modelling strategy that combines transcriptomics and phosphoproteomics datasets with prior knowledge annotated in public databases such as SIGNOR ^17^ and PhosphoSitePlus ^18^ (**Fig. 1A, panel c**). The strategy, dubbed “*Signaling Profiler”*, takes advantage of previously developed computational approaches ^19^, and complements them with novel additional features that allow a more exhaustive integration of phosphoproteomic data and *in silico* validation of the results (**Fig. 3A** **and Fig. S5**). Briefly:

1. We used footprint-based analysis ^19^ to infer the activity of key proteins: transcription factors (from the transcriptome data) and kinases and phosphatases (from phosphoproteomics data). In addition, we implemented a novel approach to infer the activity of phosphoproteins being target of (de)phosphorylation modifications (PhosphoSCORE) by combining the regulatory role and experimental fold-change of phosphosites.
2. We used the causal relations annotated in SIGNOR and PhosphoSitePlus, to build a naïve network connecting (i) FLT3, (ii) inferred kinases and phosphatases, (iii) their substrates and (iv) inferred transcription factors.
3. We exploited the CARNIVAL software ^20^ to derive FLT3^ITD-JMD^ and FLT3^ITD-TKD^ specific mechanistic models.
4. To *in silico* validate the results, we inferred the activity of key apoptotic markers as a proxy of the behavior of the two models.

**Figure 3.**
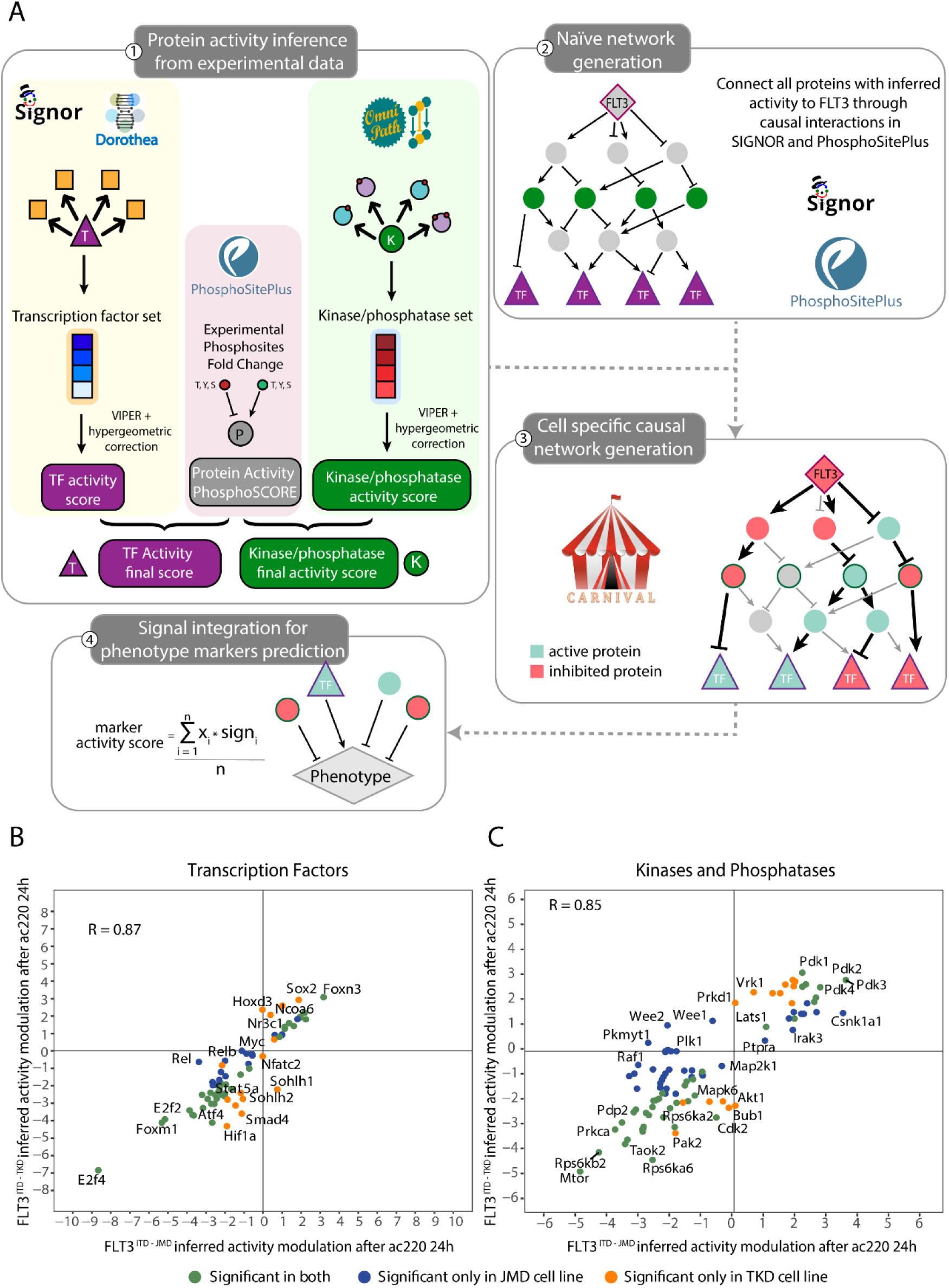
**(A)** Schematic representation of the *Signaling Profiler* workflow. **Step 1.** Protein activity of transcription factors, kinases and phosphatases was computed from experimental data using the footprint-based analysis and the ‘phosphoSCORE’ method. When needed, the two scores were averaged. **Step 2.** Proteins derived from step 1 were linked to FLT3 and to each other’s via paths of causal interactions extracted from PhosphoSitePlus and SIGNOR databases to build a naïve network. **Step 3.** CARNIVAL was used to search in the naïve network causal circuits coherent with protein activity. More specifically, in the first run we retrieved paths between FLT3 and kinases, phosphatases and substrates, whereas the second run connected all the proteins obtained from the first run with transcription factors. Eventually, the two networks were merged together. **Step 4.** The activity of protein markers of phenotypes (e.g. apoptosis) were predicted integrating the signal from upstream nodes in each cell-specific optimized network. (**B**, **C**) Protein activity prediction results. Scatterplots showing the comparison between protein activity predicted from FLT3^ITD-JMD^ (x-axis) and FLT3^ITD-TKD^ (y-axis) datasets for transcription factors (**B**) and kinases and phosphatases (**C**). Each dot represents a gene/protein, and the color indicates whether the prediction is statistically significant in both cell lines (green) or exclusively in one cell line: ITD-JMD (blue) or ITD-TKD (orange). R indicates Pearson correlation.

The first step of the *Signaling Profiler* pipeline allowed us to compute the activity of 101 kinases, 22 phosphatases and 70 transcription factors (**Fig. 3 B-C** and **Fig. S6**). As displayed in **Figure 3 C**, there is a high correlation between protein activities predicted in the two cell lines (R = 0.85 – 0.87), with a few exceptions: WEE1, WEE2 and PKMYT1 kinases are predicted to be inactive in the FLT3^ITD-JMD^ and active in the FLT3^ITD-TKD^ cells.

Protein activities of key signaling proteins are then used to feed the CARNIVAL tool together with causal networks (**Fig. 3A** **step 2 and 3**), to obtain two cell-specific models (**Fig. S7-S8**). These two graphs are static representations of the remodeling of the signal transduction cascade induced by 24 hours of quizartinib treatment. The comparison between sensitive and resistance signaling networks has the potential to reveal potential mechanisms of drug resistance and new therapeutic targets.

We next decided to monitor differences of the two FLT3-ITD specific signaling network models at a more granular level (**Fig. 3A, step 4).** Briefly, we checked whether the two models display differential modulation of pro-apoptotic and pro-survival proteins, in agreement with the phenotypes observed in the two experimental systems. As shown in **Supplementary Figure 9**, FLT3^ITD-TKD^ cells display a stronger activation of pro-survival proteins (especially, MCL1 and BCL2) and inhibition of apoptotic proteins (in particular, BAD and BIM/BCL2L11) compared to FLT3^ITD-JMD^ cells. Interestingly, CDK1 appears as a key upstream regulator of four out of five pro-apoptotic and anti-apoptotic proteins (**Fig. S9**).

### FLT3-ITD insertion site impacts the WEE1-CDK1 axis impairing cell cycle progression and apoptosis in TKIs treated cells

Given the promising role of CDK1 in mediating TKI resistance in our FLT3-TKD model, we extracted the sub-cascade that leads to its deregulation (**Fig. 4A**). As displayed in the diagram in **Figure 4A**, CDK1 is more inhibited in FLT3^ITD-TKD^ cells rather than FLT3^ITD-JMD^ cells and its regulation, downstream of FLT3, involves p27/CDKN1B, the kinase WEE1 and the phosphatase CDC25B. Interestingly, the activity of WEE1 is oppositely regulated in the two cell lines (**Fig. 3C** **and** **Fig. 4A**) and the inhibitory interaction between WEE1 and CDK1 is FLT3^ITD-TKD^ specific (**Fig. S7 and Fig. S8**). We, therefore, speculate that the WEE1-CDK1 path might play a pivotal role in inducing survival in quizartinib resistant FLT3^ITD-TKD^ cells (**Fig. 4B**).

**Figure 4.**
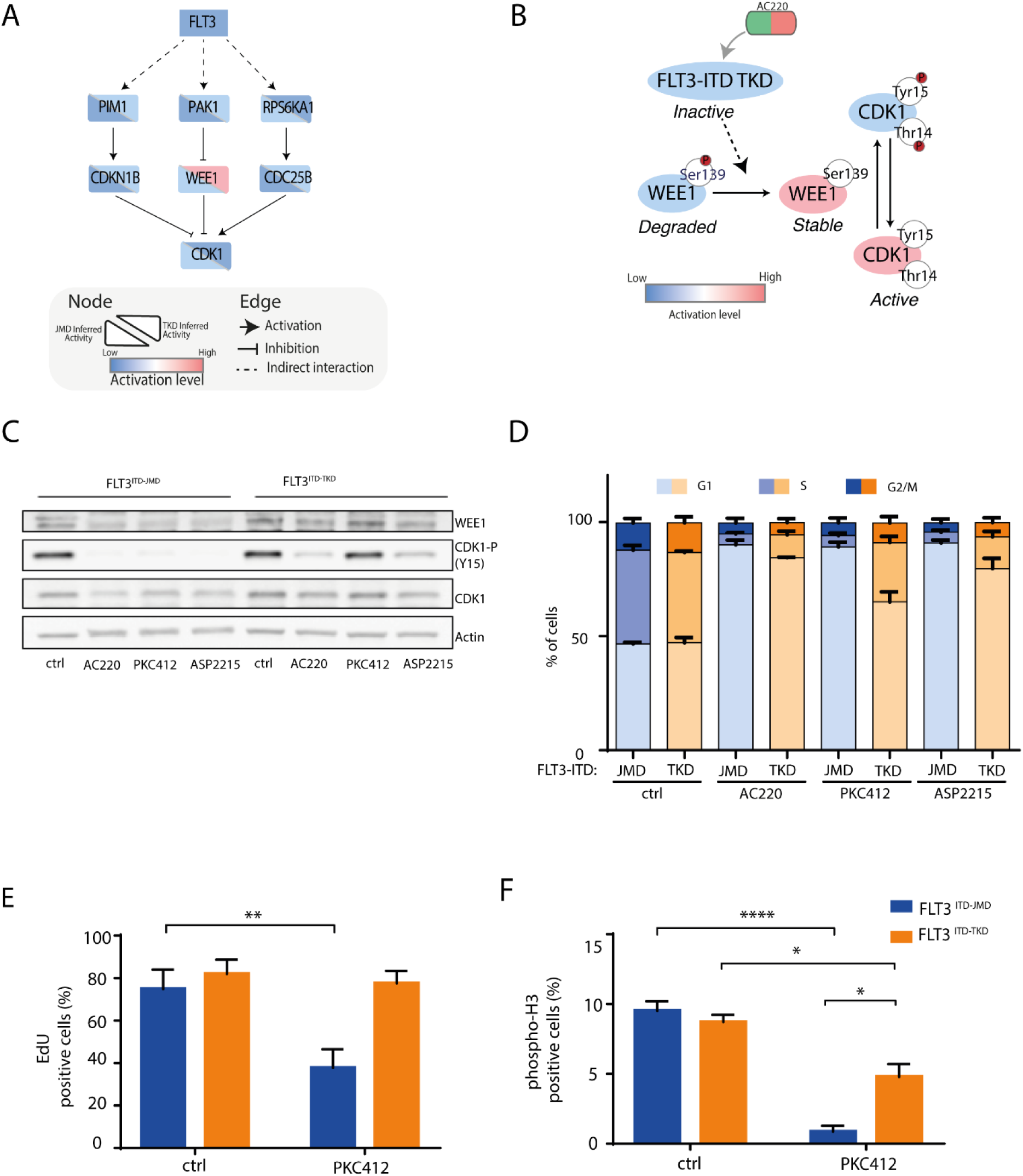
(**A**) FLT3 - CDK1 signal cascade. FLT3^ITD-TKD^ specific mechanistic model highlighting the regulation of CDK1 downstream of FLT3. For comparative analysis, for each node, the activation state in both cell line is shown (down-left half for ITD-JMD, and top-right half for ITD-TKD). Activated proteins are marked in red, whereas inhibited ones in blue. (**B**) WEE1 – CDK1 mechanistic model. Cartoon representing the potential molecular mechanism of chemoresistance suggested by the *Signaling Profiler* analysis. In FLT3^ITD-TKD^ cell line, quizartinib (AC220) -triggered inhibition of FLT3 leads to the dephosphorylation of WEE1 at Ser139, this prevents WEE1 degradation, stabilizing the protein; as such, CDK1 is phosphorylated on its inhibitory residues Tyr15 and Thr14 becoming inactive ^37^. (**C**) Representative western blot showing the phosphorylation level of CDK1 on Tyr15 and the protein level of WEE1 kinase in FLT3^ITD-JMD^ and FLT3^ITD-TKD^ BaF3 cells treated for 24 hours with 20nM quizartinib (AC220), 100nM midostaurin (PKC412) and 50nM gilteritinib (ASP2215). (**D**) Cell cycle analysis. Boxplots displaying the percentage of FLT3^ITD-JMD^ (in blue) and FLT3^ITD-TKD^ (in orange) cells in the different phases of the cell cycle as determined by flow cytometry using DAPI labeling, after treatment for 24 hours with 20nM quizartinib (AC220), 100nM midostaurin (PKC412) and 50nM gilteritinib (ASP2215). (**E**) Effect of midostaurin on cell division. FLT3^ITD-JMD^ (blue) and FLT3^ITD-TKD^ (orange) BaF3 cells were treated with 100 nM midostaurin (PKC412) for 24 hours. Percentage of cells in division was assessed by EdU labelling and flow cytometry analysis. (**F**) Bar plot representing the percentage of cells in mitosis. FLT3^ITD-JMD^ (blue) and FLT3^ITD-TKD^ (orange) BaF3 cells were treated with 100 nM PKC412 for 24 hours. Cells expressing phospho-H3 (S10) were identified by flow cytometry analysis.

WEE1 is a tyrosine kinase that is a crucial component of the G2-M cell cycle checkpoint that prevents entry into mitosis in response to cellular DNA damage by negatively regulating the activity of CDK1, the key switch responsible for M-phase initiation ^21^.

Our experiments demonstrated that TKIs treatment differently changes the abundance of WEE1 in FLT3-ITD cells (**Fig. 4C**), without affecting its transcript level (**Fig. S10A**). Consistently, in our MS-based phosphoproteomic approach, the phosphorylation level of the serine 139, which has been demonstrated to correlate with its degradation ^22^, is lower in FLT3^ITD-TKD^ cells as compared to FLT3^ITD-JMD^ cells (**Fig. S10B and Table S3**). In FLT3^ITD-TKD^ cells, the increased WEE1 protein level is associated with the enhanced phosphorylation of CDK1 (at tyrosine 15) and consequently with its negative enzymatic regulation (**Fig. 4C**). Altogether, our results provide evidence that TKI treatment, especially midostaurin (PKC412), triggers CDK1 activity in FLT3^ITD-JMD^ cells, but not in FLT3^ITD-TKD^ cells.

Thus, we tested whether the different modulation of CDK1 affects the distribution of the different phases of the cell cycle upon 24h of TKIs treatment. The cell cycle analysis revealed that the TKI-treated FLT3^ITD-JMD^ cells show a reduction of the percentage of cells in the S and G2/M phases. After exposure to TKIs, especially midostaurin (PKC412), FLT3^ITD-TKD^ cells show a higher percentage of S-phase and G2-M phase cells as compared to FLT3^ITD-JMD^ cells (**Fig. 4D**). Consistently, we observed that midostaurin significantly reduces cell proliferation only in FLT3^ITD-JMD^ cells, as revealed by the EdU assay (**Fig. 4E**). Accordingly, phosphoH3 staining revealed that the percentage of mitotic cells is significantly lower in FLT3^ITD-JMD^ as compared to FLT3^ITD-TKD^ cells upon midostaurin exposure (**Fig. 4F**).

Finally, our observations indicate that the different locations of the ITD in the FLT3 receptor impact crucial kinases determining distinct modulations of cell cycle progression.

### WEE1 kinase inhibition reverts the TKI-therapy resistance of FLT3^ITD-TKD^ cells

As we have demonstrated that FLT3-ITD resistant cells are characterized by an increased stability of the WEE1 kinase, we next investigated whether pharmacological inhibition of WEE1 would potentiate the pro-apoptotic effect of TKIs in FLT3^ITD-TKD^ cells.

Briefly, we treated FLT3^ITD-JMD^ and FLT3^ITD-TKD^ cells with adavosertib (MK1775), a highly selective WEE1 inhibitor ^23^, separately or in combination with midostaurin. As displayed in **Figure 5A**, adavosertib treatment efficiently reduces the level of CDK1 phosphorylation at Tyr15, showing efficient target engagement. Apoptotic and cell survival assays showed that WEE1 inhibitor synergises with midostaurin to trigger cell death of FLT3^ITD-TKD^ cells and to a lesser extent of FLT3^ITD-JMD^ cells (**Fig. 5B-C**).

**Figure 5.**
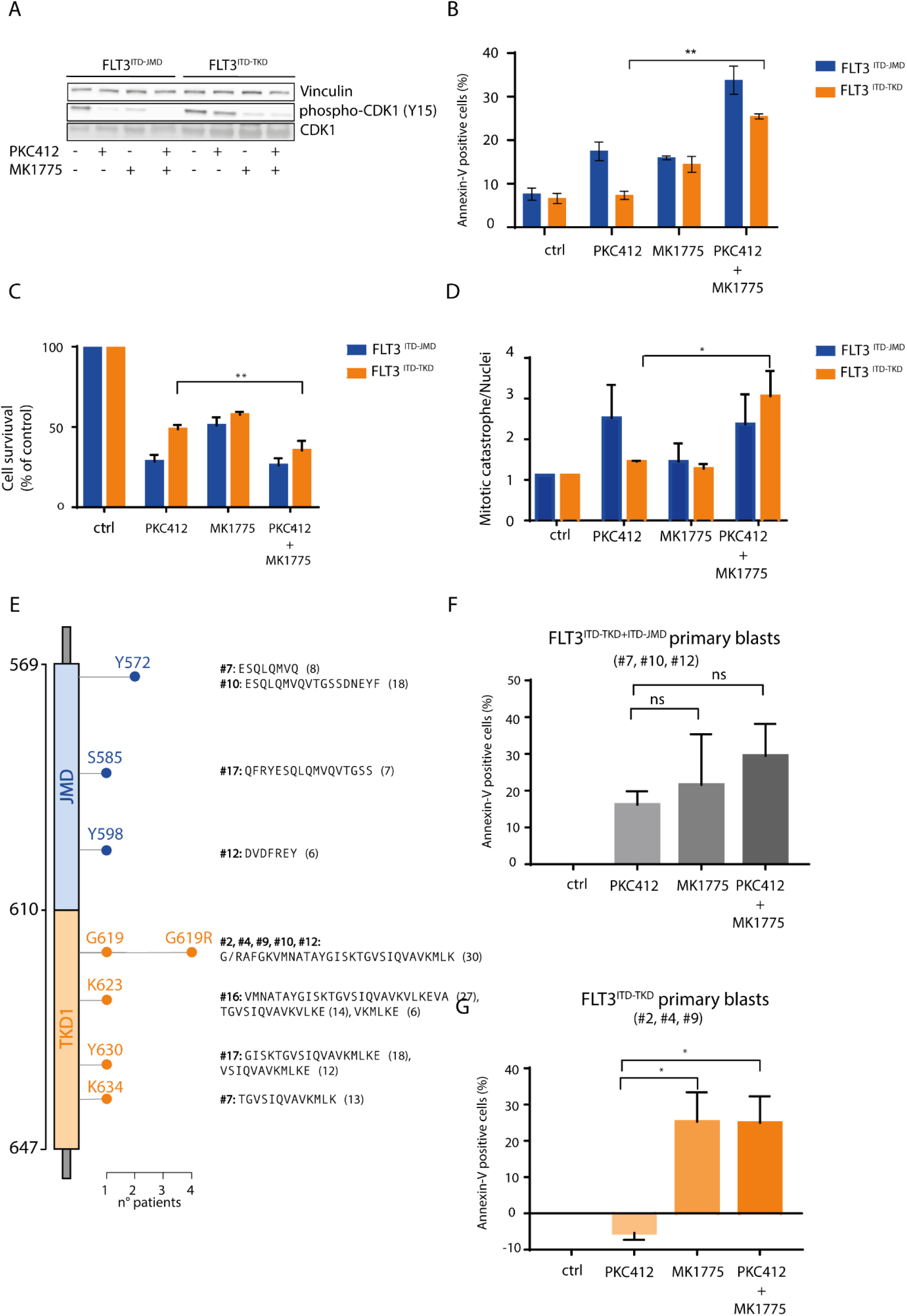
**(A)** Representative western blot showing the effect of the WEE1 inhibitor, adavosertib, on the phosphorylation levels of CDK1 on tyrosine 15. FLT3^ITD-JMD^ and FLT3^ITD-TKD^ BaF3 cells were treated with 100nM PKC412, 500nM adavosertib (MK1775) and the combination of both for 24 hours. **B**) FLT3^ITD-JMD^ (blue) and FLT3^ITD-TKD^ (orange) BaF3 cells were treated with 100nM midostaurin (PKC412), 500nM adavosertib (MK1775) and the combination of both for 24 hours. Percentage of apoptotic cells was assessed by Annexin-V labelling. (**C**) FLT3^ITD-JMD^ (blue) and FLT3^ITD-TKD^ (orange) BaF3 cells were treated with 100nM midostaurin (PKC412), 500nM adavosertib (MK1775) and the combination of both for 24 hours. Cell survival relative to control after treatment was calculated by MTT assay. **D**) FLT3^ITD-JMD^ (blue) and FLT3^ITD-TKD^ (orange) BaF3 cells were treated with 100nM midostaurin (PKC412), 500nM adavosertib (MK1775) and the combination of both for 24 hours. Percentage of mitotic catastrophe was assessed by DAPI labelling of nuclei. **(E)** Lollipop plot representing the location, amino acid sequence and length of FLT3-ITD mutations in the 9 patients analyzed. 4 patients have ITD located in TKD1 domain (#2, #4, #16, #19), 5 in both TKD1 and JMD domain (#1, #7, #10, #12, #17). Each lollipop length represents the number of patients having an ITD in that position. All mutations were derived from Sanger sequencing from primary patient blasts**. (F)** Barplot showing the percentage of treatment-induced apoptosis (*100 * (dead cells after treatment – death cells in control) / viable cells in control*) in patient-derived blasts carrying FLT3-ITD in both the JM and the TK1 domains upon the indicated treatments (patients: #7, #10, #12). Percentage of apoptotic cells was assessed by Annexin-V labelling. **(G)** Barplot showing the percentage of treatment-induced apoptosis (*100 * (dead cells after treatment – death cells in control) / viable cells in control*) in patient-derived blasts carrying FLT3-ITD exclusively in the TK1 domain upon the indicated treatments (patients: #2, #4, #19). Percentage of apoptotic cells was assessed by Annexin-V labelling.

Pharmacological inhibition of WEE1 kinase activity and the consequent removal of the G2–M checkpoint through the CDK1 hyperactivation represents an attractive strategy to drive cancer cells to enter into unscheduled mitosis and, arguably, to undergo cell death via alternative mechanisms such as the mitotic catastrophe ^24^. In line with this hypothesis, the combined treatment of midostaurin and adavosertib, exclusively, triggers mitotic cell death in FLT3^ITD-TKD^ cells (**Fig. 5D**).

We next investigated whether FLT3-ITD primary blasts, derived from 9 patients with de novo AML diagnosis, could benefit from the combined treatment of midostaurin and WEE1 inhibitor. First, blasts were isolated from peripheral blood of ITD-positive AML patients (**Fig. S11**). As expected, the molecular landscape of FLT3-ITD mutation is heterogeneous (**Fig. 5E**). We excluded one patient carrying an atypical insertion sequence in the JMD domain. Then, we classified our cohort of FLT3-ITD patients in two main groups according to the ITD insertion site (**Fig. 5E** **and Fig. S12**). This approach enabled us to obtain two subgroups: 4 FLT3^ITD-TKD^ patients (carrying the ITD in the TKD domain), 4 FLT3^ITD-JMD+ITD-TKD^ patients (carrying the ITD in both the JMD and TKD domain). We considered only patients with a single insertion, reaching three patient per subgroup. Unexpectedly, in our cohort, no patient carrying the ITD only in the JMD domain was found. Remarkably, genetic stratification based on the ITD localization reflected the drug-response phenotype (**Fig. S12**): midostaurin alone significantly triggers apoptosis in FLT3^ITD-JMD+ITD-TKD^ blasts (**Fig. 5F**), while no beneficial effects were observed in FLT3^ITD-TKD^ blasts (**Fig. 5G**). This result suggests that the pro-apoptotic effect of the ITD insertion within the JM domain is dominant over the ITD-TKD counterpart. Remarkably, both the pharmacological inhibition of WEE1 alone and the combined treatment with midostaurin trigger apoptosis of FLT3^ITD-TKD^ positive blasts, restoring their sensitivity to TKI therapy (**Fig. 5G**).

Our results provide novel evidence that the WEE1-CDK1 axis represents a promising therapeutic target to revert drug resistance in patients carrying the ITD mutation in the TKD of FLT3 that currently cannot benefit from midostaurin treatment.

## Discussion

Internal tandem duplication (ITD) in the FLT3 receptor are genetic alterations occurring in about 30% of patients with a de novo AML diagnosis. These mutations result in increased and uncontrolled kinase activity and are generally associated with poor prognostic outcomes. Although the molecular landscape of ITD mutation has been shown to be highly complex and heterogeneous, FLT3-ITD positive patients receive the same treatment consisting of standard chemotherapy combined with the multikinase inhibitor midostaurin (Richard M. Stone et al., 2018). Very recently, a retrospective explorative analysis revealed the negative prognostic impact of the ITD insertion in the tyrosine kinase domain compared to the ITD located in the juxtamembrane domain ^6^. We and others showed that the insertion site of the ITD mutations significantly impacts the ability of FLT3 inhibitors, including midostaurin, to trigger cell death in both cell lines and primary blasts ^7,9,26–28^. Consistently with previous observations, here we show that the beneficial effect of TKIs is restricted to FLT3-ITDs located in the juxtamembrane domain (JMD), but not to FLT3-ITD in the TKD region. Indeed, ITDs in the TKD alone predispose to chemoresistance and relapse, demanding for more effective and targeted treatments.

We have reported here the first unbiased, large-scale, multi-layered analysis aimed at describing the molecular mechanisms underlying the different sensitivity to TKI therapy of cells carrying FLT3-ITD mutations in the TKD or JMD domains. The main objective of this study is the identification of new potential therapeutic targets increasing the efficacy of the TKI therapy in FLT3^ITD-TKD^ patients.

Our quantitative transcriptome, proteome and phosphoproteome analysis provide an integrated picture of the TKIs-dependent molecular events. We speculate that a complex rewiring of signaling pathways may be the cause of the different sensitivity of FLT3-ITD cells to TKIs treatments. To address this point, we implemented a computational pipeline dubbed *Signaling Profiler*, that integrates readouts of transcriptome and phosphoproteome studies, with prior evidence annotated in public repositories to produce cell-specific networks representing the remodeling of signal transduction cascade at the PTM-resolution level induced by quizartinib. The observed inhibition of canonical pathways immediately downstream of FLT3 ^29^ as well as the presence of well-characterized gene products whose mutation are involved in AML progression or relapse (e.g. NPM1, CEBPA, KRAS, PTPN11) ^30^, confirm the clinical relevance of our models.

Although grounding on previously developed tools such as CARNIVAL ^20^ and VIPER ^31^, Signaling Profiler incorporates novel features such as the PhosphoSCORE calculation method, enabling for a comprehensive integration of the phosphoproteomic data; and the simulation of phenotypic biomarker, which provide the *in-silico* validation of the results.

This approach revealed a novel mechanism of resistance relying on the different regulation of the WEE1-CDK1 axis. In untreated FLT3-ITD cells, CDK1 was found to be at least partially inactivated by phosphorylation at Tyr15. Potential mechanisms upstream of CDK1 phosphorylation include the hyper-activation of the kinases WEE1, WEE2 and PKMYT1 as well as the inhibition of the phosphatase CDC25C ^32^. TKIs treatment, especially midostaurin, reversed this process and activated CDK1 to a greater extent in cells carrying the “classical FLT3-ITDs”, which are also more sensitive to apoptosis. To date, several reports remarked that an unscheduled, premature or sustained CDK1 activation has been associated to cell death through mitotic catastrophe or apoptosis induction ^32–35^. By contrast, TKIs treated cells carrying the insertion in the TKD domain are characterized by a high expression of the WEE1 kinase, which in turn determines CDK1 inactivation. Consistently, increased level of WEE1 has been shown to correlate with tumor progression and poor disease-free survival ^36^. Here, we demonstrated that deregulation of the WEE1-CDK1 axis represents a crucial mechanism of resistance to TKI therapy in FLT3-ITD positive cells and patient-derived primary blasts. Remarkably, the ability of FLT3^ITD-TKD^ cells to undergo apoptosis in response to TKI therapy was completely rescued by pharmacological inhibition of WEE1. This observation provides support for WEE1 inhibitors to be used in combination therapies with TKIs to improve the clinical outcomes of FLT3^ITD-TKD^ patients.

Interestingly, although we analyzed a small cohort of FLT3-ITD patient-derived primary blasts, our analysis confirms that insertions in the TKD sole have a significantly inferior clinical outcomes compared to blasts with insertion sites in both the JM and the TK domains (Rucker et al., 2022).

In conclusion, the results of this work endorse the discrepancies in the current practice toward the treatment of FLT3-ITD positive patients affected by AML and open-up opportunities for additional, more effective and patient-specific therapeutic strategies. Here, we speculate that these pre-clinical results create the basis of new trials that might change the clinical reality for AML patients. We suggest that FLT3-ITD patients, at diagnosis, should be stratified according to the ITD insertion site into prognostically relevant FLT3-ITD subgroups. Midostaurin maintenance therapy should be critically evaluated in case of FLT3-ITD located within the TKD1 domain and synergistic combination therapies should be used to rationally manipulate the WEE1-CDK1 axis, triggering cell death, through mechanisms that are yet to be defined. Finally, we’d like to stress the importance of unbiased, system-level studies to accelerate the investigation of more granular, patient-specific mechanisms of disease and chemoresistance toward the identification of more effective therapeutic targets.

## Acknowledgments

We thank Prof. Gianni Cesareni and Prof. Luisa Castagnoli for their essential scientific input. Dr. Serena Paoluzi and Dr. Marta Iannuccelli for their technical support. We thank Dr. Grumati and the MS facility at TIGEM for their scientific and technical support. This research was funded by the Italian association for cancer research (AIRC) with a grant to F.S. (Start-Up Grant n. 21815). V.V. is supported by MUR, G.M.P. and V.B. are supported by the AIRC Start up grant number 21815.

## Author contributions

Conceptualization, G.M., V.V., L.P., F.S.; methodology, G.M., V.V., G.M.P., N.K., F.K., D.M., M.B., T.F., L.P., F.S.; formal analysis, G.M., V.V., V.B., S.L., G.M.P., F.K., D.M., M.B.; investigation, G.M., V.V., with the contribution of F.S., L.P., V.B., S.L., G.M.P.; writing—original draft preparation, F.S., L.P.; resources, F.S.; writing—review and editing, G.M., V.V, T.F., F.S., L.P.; supervision, F.S.; funding acquisition, F.S. All authors have read and agreed to the published version of the manuscript.

## STAR Methods

### Cell culture

Mouse Ba/F3 cells expressing ITD-JMD and ITD-TKD constructs were provided by courtesy of T. Fischer. The cells were cultured in RPMI 1640 medium (Hyclone, Thermo Scientific, Waltham, MA) supplemented with 10% heat-inactivated fetal bovine serum (Euroclone ECS0090D), 100 U/ml penicillin and 100 mg/ml streptomycin (Gibco 15140122), 1 mM sodium pyruvate (Sigma-Aldrich, St. Louis, Missouri, United States, S8636) and 10 mM 4-(2-hydroxyethyl)-1-piperazineethanesulfonic acid (HEPES) (Sigma H0887). Ba/F3 cells were maintained at a density of 300.000 cells/ml in T75 flasks with vented-filter cap (Sarsedt, 50809261).

### Immunoblot analysis

BaF3 cells were seeded at a concentration of 500.000 cells/ml and treated as indicated. After treatments cells were centrifuged and washed in PBS 1x. Next, cells were lysed in ice-cold lysis buffer (150 mM NaCl, 50 mM Tris–HCl, pH 7.5, 1% Nonidet P-40, 1 mM EGTA, 5 mM MgCl_2_, and 0.1% SDS) supplemented with 1 mM PMSF, 1 mM ortovanadate, 1 mM NaF, protease inhibitor mixture 1×, inhibitor phosphatase mixture II 1×, and inhibitor phosphatase mixture III 1× and incubated for 30 min. Protein lysates were separated at 13,000*g* for 30 min. The total protein concentration was determined using the Bradford reagent. Protein extracts were denatured and heated at 95°C for 10 min in NuPAGE LDS Sample Buffer that contained DTT as a reducing agent (NuPAGE Sample Reducing Agent). Proteins were resolved using 4– 15% Bio-Rad Mini-PROTEAN TGX/CRITERION polyacrylamide gels. Proteins were transferred to Trans-Blot Turbo Mini Nitrocellulose Membranes using a Trans-Blot Turbo Transfer System (Bio-Rad), and the nonspecific binding membranes were saturated in blocking solution (5% skimmed milk powder, 0.1% Tween 20 in 1× TBS) at room temperature for 1 hour. Saturated membranes were incubated overnight with primary antibodies diluted in BSA 5% (anti-phospho FLT3 1:1000, CST 3464S; anti-phosphoSTAT5 1:1000, Abcam ab32364; anti-phosphoERK1/2 1:1000, CST 9101; anti-FLT3 1:1000, CST 3462; anti-STAT5 1:1000, Abcam ab16276; anti-ERK1/2 1:1000, CST 4695; anti-phospho CDK1 (Y15), CST 9111S; anti-CDK1/2 1:1000, Santa Cruz sc. 53219; anti-Wee1 1:1000, Abcam ab273016; anti-actin 1:3000, Sigma A2066). HRP-conjugated secondary antibodies (Goat Anti-Mouse IgG (H+L)-HRP Conjugate 1:3000, BIORAD 1721011) were diluted in blocking solution and used for the detection of the primary antibodies. Chemiluminescence was detected using Clarity Western ECL Blotting Substrates (Bio-Rad) and the Las-3000 Imaging System (Fujifilm). Band densities were quantified using ImageJ and normalized to the loading control.

### Apoptosis assay

Cells were plated at a concentration of 500.00 cells/ml and treated as indicated. After incubation for 24 hours, apoptotic cells were measured by flow cytometry using Ebioscience™ Annexin V Apoptosis Detection Kit APC according to the kit instruction (Cat. 88-8007-74, Thermo Fisher Scientific). Cells positive for annexin-V were counted as apoptotic cells.

### MTT assay

Cell viability was measured using the Cell Proliferation Kit I (MTT) (Roche). Cells were treated as indicated for 20 hours. Then, MTT was added to the cells and incubated for 4 hours at 37 ◦C. Solubilization Solution was used to dissolve the formazan crystals during an overnight incubation. Finally, the plates were read at 590nm using a microplate reader (Bio-Rad).

### Sample preparation for proteomic and phosphoproteomic analysis

Cell lysis was performed by adding SDC lysis buffer containing 4% (w/v) SDC, 100 mM Tris-HCl (pH 8.5). We used the inStageTip (iST) method for the proteome preparation ^38^. Phosphoproteome preparation was performed by the EasyPhos workflow as previously described ^39^. Briefly, per condition a total of 1 mg protein input material was lysed, alkylated and reduced in a single step. After the protein’s digestion, phosphopeptides were enriched using TiO2 beads.

### Mass spectrometry analyses

The peptides and the phosphopeptides were desalted on StageTips and separated on a reverse phase column (50 cm, packed in-house with 1.9-mm C18-Reprosil-AQ Pur reversed-phase beads) (Dr Maisch GmbH) over 120 min or 140 min (single-run proteome and phosphoproteome analysis respectively). After elution, peptides were electrosprayed and analyzed by tandem mass spectrometry on a Orbitrap Exploris 480 (Thermo Fischer Scientific). The instrument was set to alternate between a full scan followed by multiple HCD based fragmentations scans for a total cycle time of up to 1 s.

### RNAseq analysis

BaF3 cells were cultured in growth medium at the density of 500.000 cell/ml. Next, AC220 was added at a concentration of 20nM and incubated for 24 hours. Then, cells were centrifuged at a speed of 300 x g, washed in PBS and total RNA was isolated from the harvested cells using RNeasy micro kit (Qiagen, Hilden, Germany 74004) and was quantified using the Qubit 2.0 fluorimetric Assay (Thermo Fisher Scientific). Libraries were prepared from 100 ng of total RNA using the QuantSeq 3’ mRNA-Seq Library Prep Kit FWD for Illumina (Lexogen GmbH, Vienna, Austria) and their qualities were assessed by using screen tape High sentisivity DNA D1000 (Agilent Technologies, Santa Clara, California, United States). Libraries were sequenced on a NovaSeq 6 000 sequencing system using an S1, 100 cycles flow cell (Illumina Inc., San Diego, California, United States). Illumina novaSeq base call (BCL) files were converted into fastq file by bcl2fastq (version v2.20.0.422). Sequence reads were trimmed using bbduk software (bbmap suite 37.31) in order to remove adapter sequences, poly-A tails and low-quality end bases (regions with average quality below 6). Alignment was performed with STAR 2.6.0a ^40^ on mm10 reference assembly obtained from cellRanger website (Ensembl assembly release 93). The expression levels of genes were determined with htseq-count 0.9.15 (Ref. 29) by using mm10 Ensembl assembly (release 93) downloaded from cellRanger website.

### Proteome and Phosphoproteome Data processing

Raw mass spectrometry data were analyzed in the MaxQuant environment ^41^, version 1.5.1.6, employing the Andromeda engine for database search. Proteome and phosphoproteome samples were analysed together by specifying two separate groups and setting group specific parameters for each sample type. MS/MS spectra were matched against the *Mus musculus* UniProtKB FASTA database (September 2014), with an FDR of < 1% at the level of proteins, peptides and modifications. Enzyme specificity was set to trypsin, allowing for cleavage N-terminal to proline and between aspartic acid and proline. The search included cysteine carbamidomethylation as a fixed modification. Variable modifications were set to N-terminal protein acetylation and oxidation of methionine as well as phosphorylation of serine, threonine tyrosine residue (STY) for the phosphoprotemic samples. MaxQuants Label free Quantification method and a minimum ration count of two was used for the total proteome samples. For proteome and phosphoproteome analysis, where possible, the identity of peptides present but not sequenced in a given run was obtained by transferring identifications across liquid chromatography (LC)-MS runs (‘match between runs’). For phosphopeptide identification, an Andromeda minimum score and minimum delta score threshold of 40 and 17 were used, respectively. Peptides had to be fully tryptic in both proteome or phosphoproteme samples and up to two or four missed cleavages were allowed for protease digestion, respectively.

### Proteome and Phosphoproteome Bioinformatics Data Analysis

Bioinformatic analysis was performed in the Perseus software environment ^42^. Statistical analysis of proteome and phosphoproteome were performed on logarithmized intensities for those values that were found to be quantified in any experimental condition. Phosphopeptides intensities were normalized by subtracting the median intensity of each sample. Student t-Test with a permutation-based FDR cutoff of 0.07 and S0 = 0.1 was performed to identify significantly modulated proteins and phosphopetides between two different conditions. Categorical annotation was added in Perseus in the form of GO biological process (GOBP), molecular function (GOMF), and cellular component (GOCC), KEGG pathways and kinase substrate motifs (extracted from HPRD). Concerning the kinase substrate motifs, we performed a 1D annotation enrichment analyses to identify statistically significant enriched kinase-substrates motifs in AC220 treated cells ^43^. Multiple hypothesis testing was controlled by using a Benjamini-Hochberg FDR threshold of 0.05.

### EdU incorporation assay

For the proliferation assay cells were seeded at the concentration of 500.000 cells/ml and treated for 24 hours with 100nM PKC412. During the last two hours of incubation 5-ethylnyl-2′-deoxyuridine (EdU) was added at a concentration of 10 μM. After incubation, 1×10^6^ cells were centrifuged at 300g for 5 min and then washed in PBS1X. The Click-iT® reaction to detect EdU positive cells was performed according to the manufacturer’s instructions. Percentage of cell in division was assessed by flow cytometry.

### Phospho-H3 labeling

Celle were seeded at a concentration of 500.00 cells/ml and treated with 100nM PKC412 for 24 hours. After treatment, 1×10^6^ cells were centrifuged at 300g for 5 min and then washed once in PBS1X. Cells were fixed with 70% ethanol overnight at 4°C. Next, cells were centrifuged at 300xg for 10 min and washed once with PBS1X+2%BSA. 500µl of PBS1X+1% saponin was added and incubated for 15 min at room temperature to permeabilize cells. After a wash with PBS1X+2%BSA, cells were incubated with the anti-phospho-H3 antibody (Abcam AB267372) diluted 1:500 in PBS1X+1%BSA+0.5% saponin for 90 min at room temperature. After a wash in PBS1X+1%BSA, cells were incubated with 100ul of an anti-rabbit Alexa Fluor 455 diluted 1:200 in PBS1X+1%BSA+0.5%saponin for 1h at room temperature. After the incubation, cells were washed with PBS1X twice. The percentage of phosphor-H3 positive cells were quantified by flow cytometry.

### Primary patient blast analyses

Peripheral blood (PB) samples from AML patients were obtained upon patient’s informed consent and in accordance with the declaration of Helsinki (ethics committee approval number: 115/08). The integration site of the FLT3-ITD mutation was determined as previously described ^44^. Briefly, RNA was prepared from PBMCs using the RNeasy Mini Kit (Qiagen, Germany), reverse-transcribed to cDNA using the SuperScript reverse transcriptase system (ThermoFisher Scientific), and the FLT3-ITD region was amplified by PCR (fw-primer: GCAATTTAGGTATGAAAGCCAGC, rev-primer: CTTTCAGCATTTTGACGGCAACC). PCR products were re-purified using the QIAquick PCR purification kit (Qiagen) and subjected to Sanger sequencing (using the same fw-primer) by Eurofins (Luxembourg).

To classify patients according to the ITD localization, we first translated the raw FASTA files obtained from Sanger sequencing using all the possible reading frames using R package “Biostrings” ^45^. Using blastp ^46^, we then aligned all the translated sequences to the JMD and TKD domain sequences of canonical FLT3, as annotated in UniProtKB. Manual evaluation of the alignment allowed us to identify the insertion site and duplicated sequences, as displayed in **Figure 5E**.

Mononuclear cells from the PB were obtained using Ficoll-Paque (GE Healthcare, Chicago, IL). Cryoconserved PBMCs from 12 patients were cultured at a density of 5×10^5^ /mL in RPMI-1640 (Sigma-Aldrich, St. Louis, MO) supplemented with 10% FCS (c.c.pro, Germany), 2 mM L-glutamine (Sigma-Aldrich), and 40 U/mL Penicillin-Streptomycin (ThermoFisher Scientific) for 24h in absence or presence of PKC412, MK1775 or a combination of both.

Viability of the AML blasts was determined by flow cytometry using Annexin V – APC and 7AAD together with the Annexin V staining buffer according to the manufacturers’ instruction (Biolegend, San Diego, CA). Prior to viability staining, samples were stained with fluorochrome-coupled antibodies after blocking with human IgG (Gamunex, Grifols, Barcelona, Spain). Samples were recorded on a Cytek NL-3000 spectral flow cytometer. First, blasts were gated based on the CD45/SSC distribution and on FSC-A/FSC-H parameters to identify single cells population. Then, blasts were gated according to the expression of typical blast markers (CD33, CD34, CD13, CD117). Data was analyzed using FlowJo V10 (Becton-Dickinson, Franklin Lakes, NJ).

### Statistics

All the experiments have been conducted in at least 3 independent replicates obtained from 3 cell line batches (n = 3). Data are presented as means ± standard error of the mean (SEM). Multiple comparisons between three or more groups were performed using one-way or two-way ANOVA. Statistical significance between two groups was estimated using the unpaired t test assuming a two-tailed distribution. Statistical significance is defined as *p < 0.05; **p < 0.01; ***p < 0.001. All statistical analyses were performed using Prism 7 (GraphPad).

### SIGNALING PROFILER

#### Causal interaction database download

We downloaded all the causal interactions available for *Mus musculus* (TaxID = 10090) and *Homo sapiens* (TaxID = 9606) from the SIGNOR ^16^ and PhosphoSitePlus® ^47^ resources. SIGNOR 2.0 datasets were downloaded via rest API and refer to December 2021. Interactions in SIGNOR annotated with ‘down-regulates’, ‘up-regulates’ or ‘unknown’ were assigned values -1, 1 and 0, respectively.

Causal phosphorylations were extracted from PhosphoSitePlus® ^18^ by manually downloading and combining two independent tables: (i) kinase-phosphosite interactions (‘Kinase_Substrate_Dataset.gz’), (ii) regulatory role of phosphosites on protein (‘Regulatory_sites.gz’). Tables were joined using, as key, the UNIPROT ID and the modified residue. We, next, manually mapped the content of the ‘ON_FUNCTION’ column (representing regulatory role of phosphosites) into 1, -1, 0 values.

The so manipulated datasets were, then, combined together and filtered to retain interactions with a defined regulatory effect (−1 or 1).

The results of this pipeline are two causal interactomes accounting for 19,310 and 25,948 interactions in *Mus musculus* and *Homo sapiens*, respectively.

#### Protein activity prediction

##### Footprint-based analysis

Transcription Factors-target genes collection was retrieved from DoRothEA R Package (v. 1.6.0, organism *Mus musculus,* confidence: A) ^48^ and from SIGNOR (filtering for transcriptional regulations). Kinase-substrates and phosphatase-substrate collections were retrieved from Omnipath ^49^. To estimate kinases and phosphatases’ activity from substrates and transcription factors’ activity from target genes, we used the VIPER algorithm ^31^. We used as phosphosite and gene level statistic (VIPER parameters) their experimental fold change. We set the eset.filter parameter to FALSE. We included proteins with at least 1 measured transcript (or phosphosite). We retained only proteins with enrichment *p-value* < 0.05 in at least one cell line. We obtained the inferred activity of 51 transcription factors, 94 kinases and 20 phosphatases.

Hypergeometric test implemented in RVenn package (v. 1.1.0) was used to derive the p-value that a protein can be significantly enriched by chance. For each protein, we used enrichment_test function with measured genes in each regulon as *set1*, significant genes in each regulon as *set2* and all measured genes as *univ*. Log10(pvalue) scaled in [0,1] range was used to weight each protein VIPER score in order to give less importance to proteins with fewer significantly modulated targets in experimental data.

##### PhosphoSCORE

We derived the Quizartinib-induced activity modulation of proteins that are target of (de)phosphorylation from the relative phosphoproteomics data and from the regulatory role of phosphosites parsed from SIGNOR and PhosphoSitePlus (as described in the Causal interaction database section), using the formula:

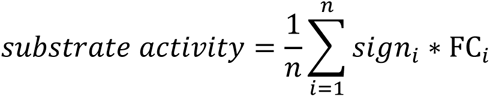

where: *n* is the number of phosphosites regulating a target, *sign_i_* is the regulatory role of phosphosite (1 or -1) and *FC*_i_ is the experimental fold change of phosphosites significantly modulated in at least one cell line.

The regulatory role of phosphosites was inferred from the mouse and the human datasets and combined by orthology mapping, using the blastp software ^46^.

This process allowed us to predict the activity of 7 additional kinases, 19 transcription factors, 2 phosphatases and 70 proteins with other molecular functions. For 3 transcription factors and 11 kinases both VIPER weighted score and PhosphoSCORE were present, in this case we averaged the two scores.

#### Cell – specific naïve causal network generation

Murine causal interactions from SIGNOR and PhosphoSitePlus were converted in a graph using ‘igraph’ R package (v. 1.2.10). We, then, extracted a subnetwork containing: (i) all the shortest paths from FLT3 to kinases, phosphatases and substrates; (ii) kinase – substrate and phosphatase-substrates interactions; (iii) all the shortest paths from kinases, phosphatases and substrates to transcription factors. We, thus, obtained a naïve network containing 871 nodes and 3422 edges.

#### CARNIVAL

For each cell line, we optimized the naïve network on inferred protein activity values using CARNIVAL R package ^20^. CARNIVAL filters the naïve network retaining causal paths coherent with the activity of start and end nodes. The naïve network was pre-processed to: (i) remove incoming edges in FLT3 and (ii) remove feedback loops. We, then, performed two runs of CARNIVAL: The first run starting from the FLT3 to kinases, phosphatases and substrates; the second from all nodes present in the output of the previous run to transcription factors. FLT3 activity was assigned a -1 value, since quizartinib (AC220) inhibits its activity, whereas the activity of the remaining nodes was assigned as described in the *Protein activity prediction* section.

The networks obtained from the two runs were merged to generate two final, cell-specific networks linking FLT3 to transcription factors, namely the FLT3^ITD-JMD^ model (210 nodes and 363 edges) and the FLT3^ITD-TKD^ model (201 nodes and 322 edges).

### Text mining approach

We used a text-mining approach to query the literature database, Europe PMC, to identify research articles characterizing pro-apoptotic and pro-survival proteins associated to FLT3-ITD AML. The query is: (“pro-apoptotic” OR “anti-apoptotic” OR “pro-survival” OR “anti-survival”) AND (TITLE:“FLT3-ITD” AND (TITLE:“AML” OR TITLE:“Acute Myeloid Leukemia”))AND (OPEN_ACCESS:y) AND (PUB_TYPE:“Research-article” OR PUB_TYPE:“report”). We derived a list of eleven pro- or anti-apoptotic proteins (Table S5).

### Phenotype marker prediction

To *in silico* validate the results, we extracted from SIGNOR and PhosphoSitePlus direct (one step) connections between nodes in the optimized models and phenotype markers derived from the text mining approach. We were able to connect only five out of eleven proteins, since (i) some proteins were already in the network (e.g. MYC); (ii) some biomarkers didn’t have any direct link (e.g. PIM1, XIAP, BCL2L10); (iii) some proteins displayed contradictory literature evidence (e.g. PARP1, BIRC5).

We integrated the signal on each marker to derive its activity modulation after quizartinib. The impact of each regulator over the apoptotic marker was computed multiplying its activation state, as inferred from the experimental data (as described in the *Protein activity prediction* section), by the sign of regulation, namely -1 for inhibitions and +1 for activations. Finally, all the effects on each marker were averaged to derive the activity score.

## Supplementary material

**Figure S1.**
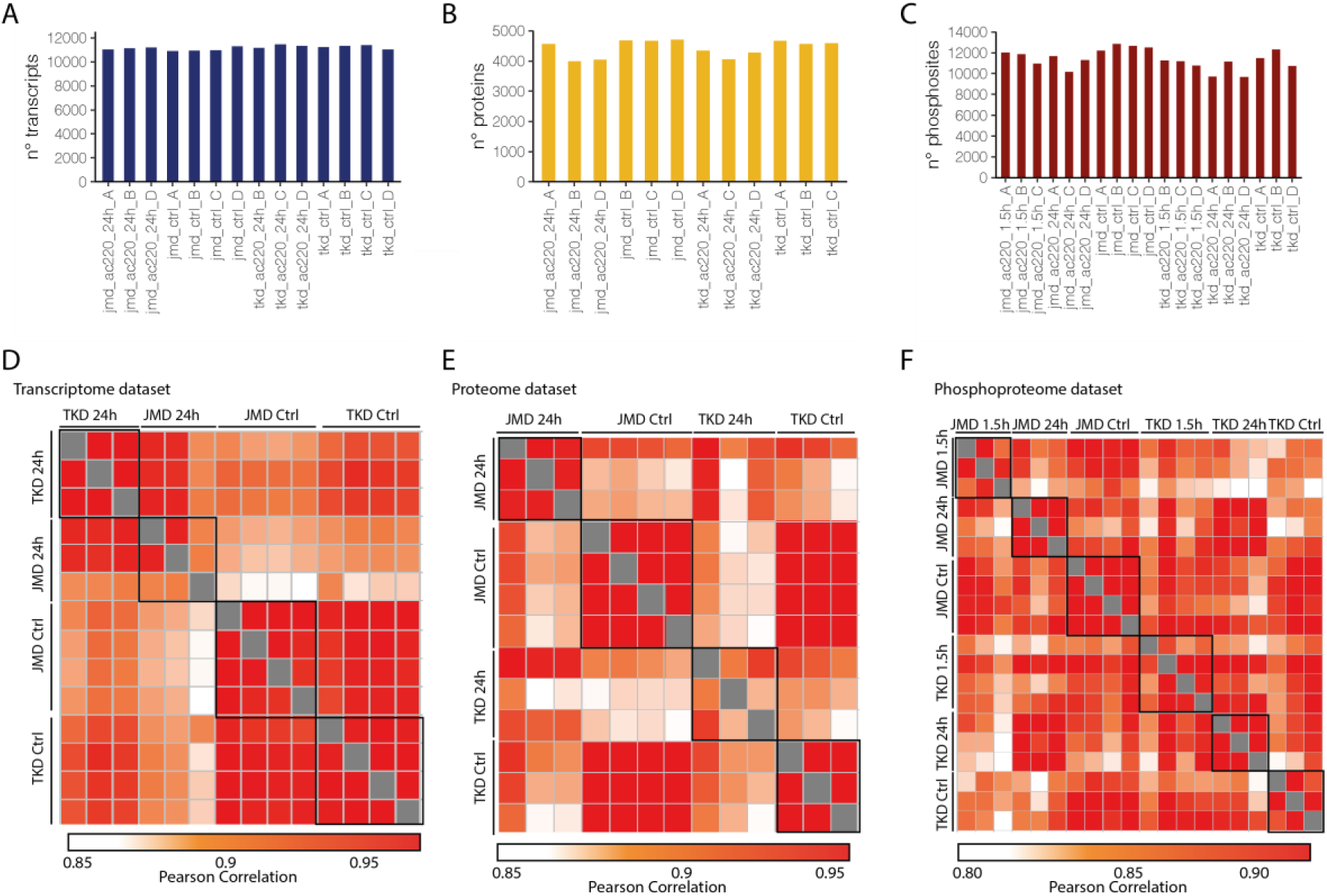
High coverage and reproducibility of proteome and phosphoproteome data. (**A-C**) Number of quantified transcripts (**A**), proteins (**B**) and phosphosites (**C**) in biological replicates of the indicated experimental conditions. (**D-F**) Heatmap showing the Pearson correlation coefficients between the different biological replicates in the trascriptome (**D**), proteome (**E**) and phosphoproteome (**F**) datasets.

**Figure S2.**
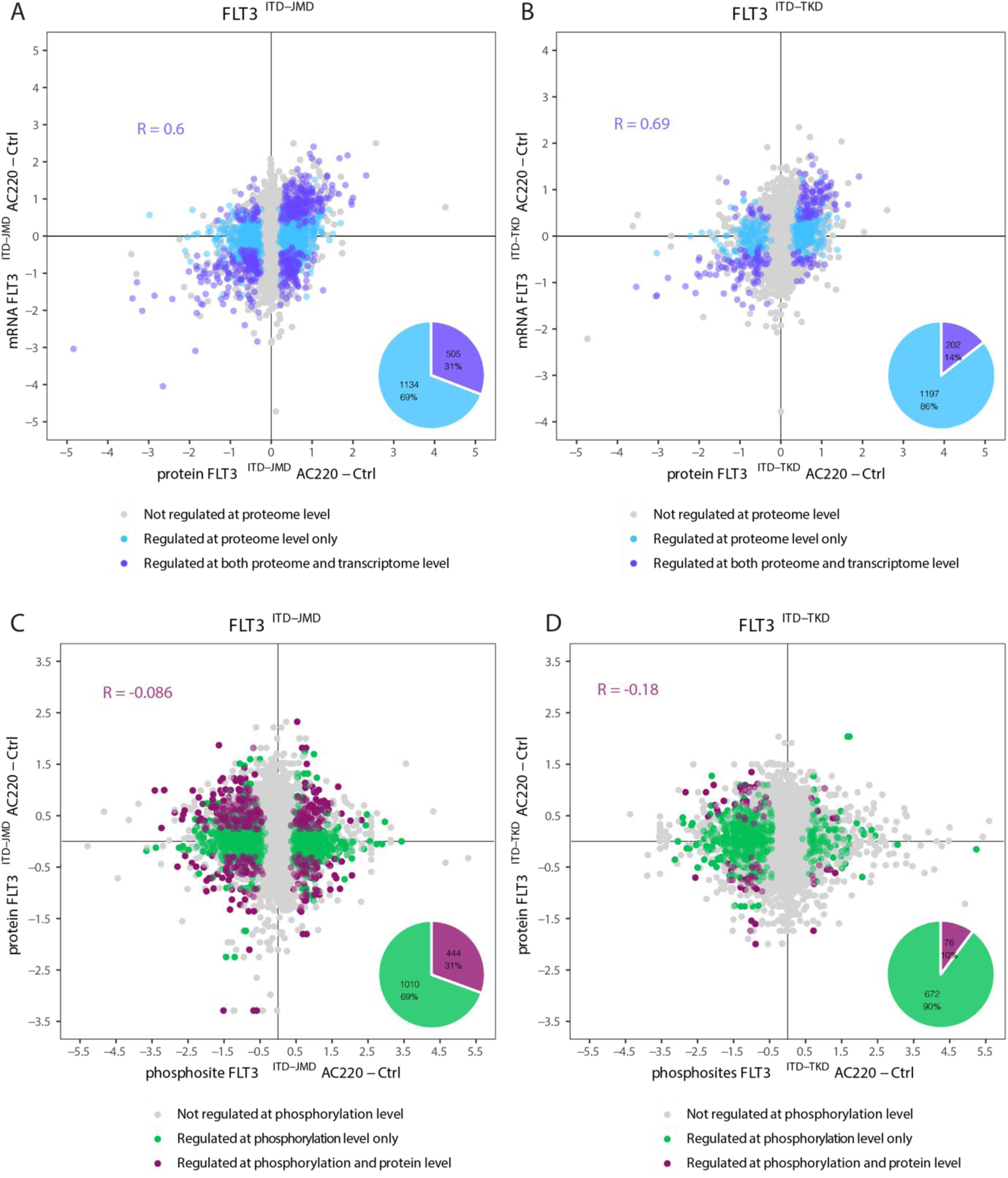
Transcriptome, proteome and phosphoproteome comparative analysis. **A-B)** Correlation analysis between protein and mRNA levels in FLT3^ITD-JMD^ cells (**A**) and FLT3^ITD-TKD^ cells (**B**). Proteins (dots) significantly modulated both at proteome and trascriptome level are marked in violet, whereas those modulated exclusively at the proteome level are indicated in blue. The pie chart details corresponding percentages. **C-D)** Correlation analysis between protein and phosphorylation levels in FLT3^ITD-JMD^ cells (**C**) and FLT3^ITD-TKD^ cells (**D**). Phosphosites (dots) modulated by quizartinib (AC220) treatment and belonging to proteins modulated in quantity (proteome level) are represented in purple, whereas those modulated only at the phosphorylation level are marked in green.

**Figure S3.**
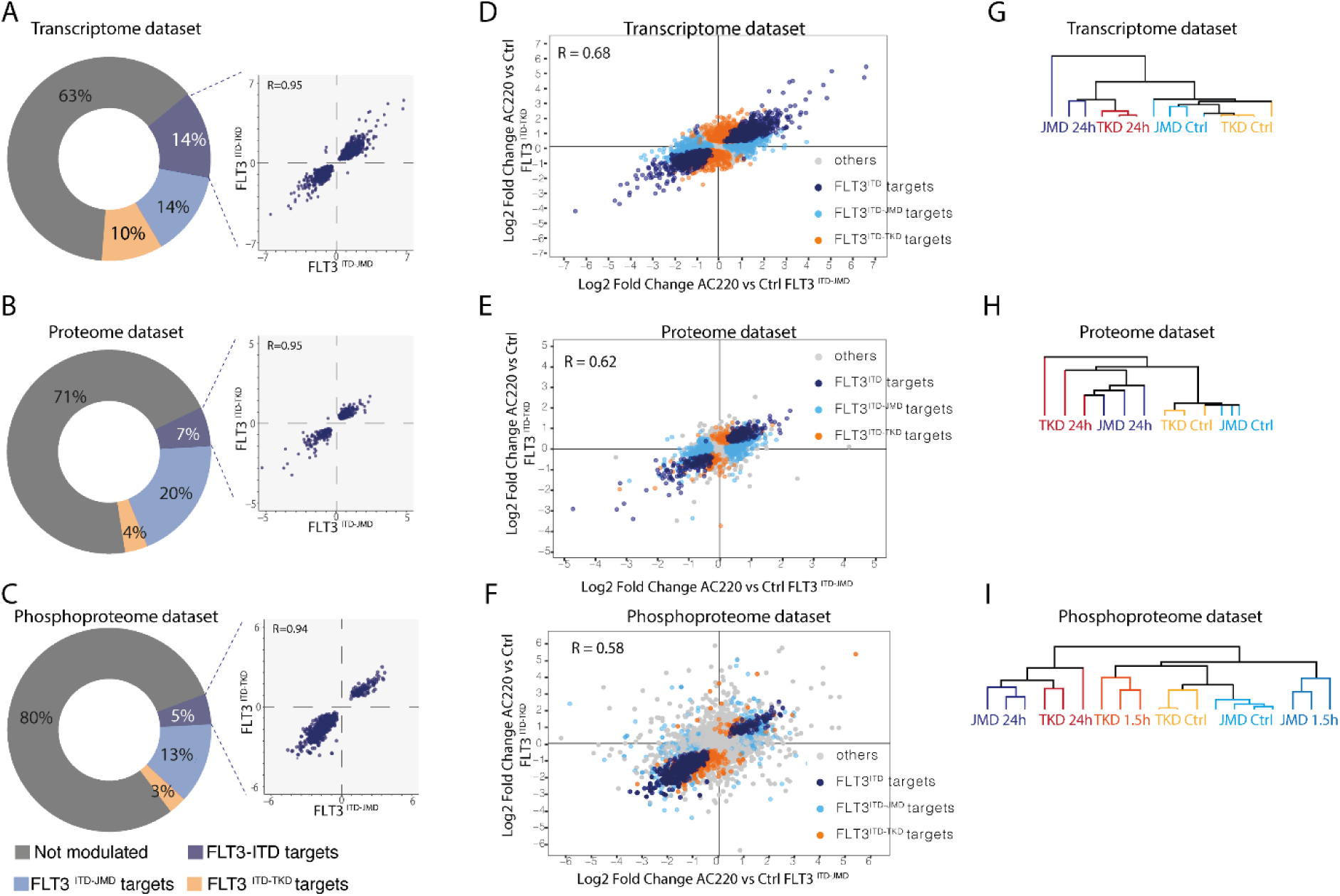
Comparative analysis of quizartinib-induced changes at the transcriptome, proteome and phosphoproteome levels in FLT3^ITD-JMD^ and FLT3^ITD-TKD^ cells. **(A-C)** Donut charts indicating the percentage of quizartinib significantly modulated transcripts (**A**), proteins (**B**) and phosphosites (**C**) in FLT3^ITD-JMD^ and/or FLT3^ITD-TKD^ cells. For the analytes modulated in FLT3^ITD-JMD^ and FLT3^ITD-TKD^, the comparison of transcript, protein and phosphosite level is shown in a scatterplot with Pearson Correlation. **(E-G)** Comparison of mRNA (**E**), protein (**F**) and phosphosites (**G**) fold change between quizartinib (AC220)-treated FLT3^ITD-JMD^ (x-axis) and FLT3^ITD-TKD^ (y-axis) cell lines. Each dot corresponds to a transcript, protein or phosphosite quantified in both cell lines. Analytes can be significantly modulated by quizartinib (AC220) in both cell lines (blue), or exclusively in one cell line: FLT3^ITD-JMD^ (light blue) or FLT3^ITD-TKD^ (orange). Global Pearson correlation is shown. (**G-H-I**) Unsupervised hierarchical clustering (Pearson correlation distance) of the log2 intensity of more than 11,000 transcripts (**G**), 5,000 proteins (**H**) and 16,000 phosphosites (**I**), as indicated.

**Figure S4.**
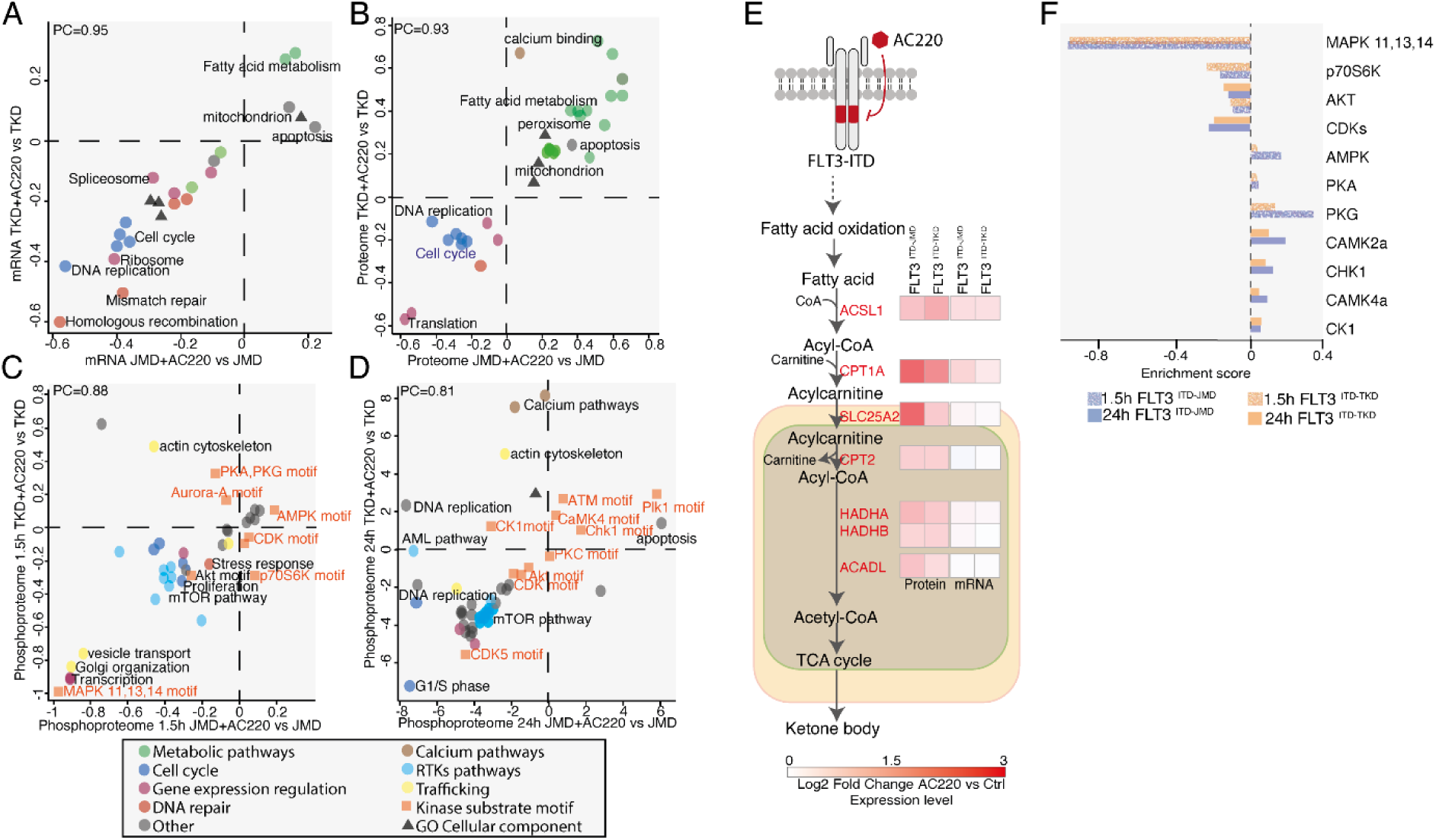
Global pathways modulation in quizartinib treated FLT3^ITD^ cells. Two-dimensional annotation enrichment analysis. Pathways modulated in quizartinib treated FLT3^ITD-TKD^ cells (y-axis) at the transcriptome (**A**), proteome (**B**), phosphoproteome (**C-D**) level in comparison with quizartinib treated FLT3^ITD-JMD^ cells (x-axis) (Benjamin Hochberg FDR < 0.05). Each dot represents a specific KEGG pathway or GO-Biological Process (GO-BP) term. Groups of related pathways or GO-BP are labeled with the same color, as described in the inset. Position scores of the pathways at the transcriptome and proteome level are indicated in the x and y axes, respectively ^43^. Negative values indicate downregulation, whereas positive values upregulation. (**E**) Schematic representation of the fatty acids oxidation; for each enzyme the corresponding quizartinib-induced change in mRNA and protein concentration in FLT3^ITD-JMD^ and FLT3^ITD-TKD^ cells is shown. (**F**) Kinase overrepresented in significantly modulated phosphosites identified in FLT3^ITD-JMD^ (blue) and FLT3^ITD-TKD^ (orange) cells.

**Figure S5.**
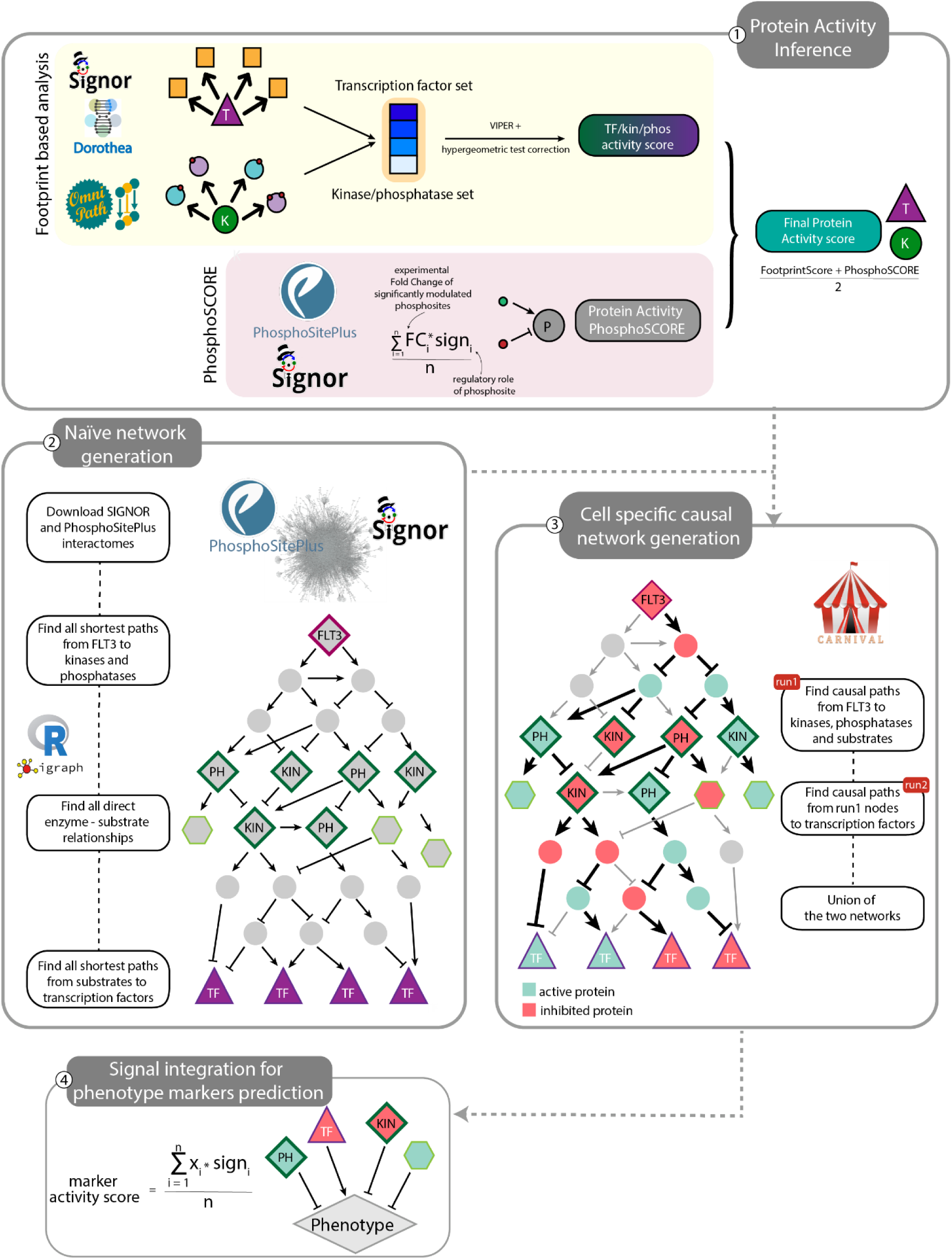
Mechanistic cell – specific causal model construction workflow. Detailed description of the *Signaling Profiler* pipeline described in Figure 4. **Step 1.** Protein activity of transcription factors (purple triangle), kinases and phosphatases (green circles) was inferred from experimental data combining the footprint-based analysis and the ‘phosphoSCORE’ technique. In the first case, protein activity is derived from the modulation of its downstream targets using the VIPER statistical tool ^31^ transcription factors’ activity is calculated from transcriptomic data whereas kinases and phosphatases’ activity from phosphoproteomic data. VIPER output is corrected through a hypergeometric test. In the second case, our de novo developed PhosphoSCORE technique exploits the modulated phosphosites of a protein and their regulatory role (as extracted from SIGNOR ^16^ and PhosphoSitePlus repositories^47^ to compute its activity (PhosphoSCORE). When a protein is assigned to both scores, they are averaged to obtain a final protein activity score. **Step 2.** Assembly of a network of causal interactions linking FLT3 and proteins characterized in step 1, to build a naïve network. Briefly, we exploited causal interactions annotated in the SIGNOR and PhosphoSitePlus resources to extract all the shortest paths from FLT3 receptor (purple diamond) to kinases and phosphatases (dark green-bordered diamonds). Similarly, we retrieved direct relationships between kinases/phosphatases and their substrates (light green-bordered hexagon). Finally, we searched for the shortest paths from kinases, phosphatases and substrates to transcription factors (purple-bordered triangle). In step 2 we considered exclusively directionality and distance and not the regulatory role of each interaction (up/down regulation). **Step 3.** CARNIVAL ^20^ was used to search in the naïve network circuits coherent with protein activity inferred in Step 1 (bold black edges): mint green nodes are predicted up-regulated after quizartinib (AC220) treatment, whereas red nodes are down-regulated. Because quizartinib (AC220) causes FLT3 inhibition, the receptor activity was set to -1 (repressed). We executed two runs of CARNIVAL: run 1 retrieved coherent paths linking FLT3 to kinases, phosphatases and substrates, run 2 linking proteins derived from run 1 with transcription factors. The two networks were, eventually, joint to obtain, for each cell line, a final mechanistic model recapitulating the signal cascade downstream of FLT3, rewired by quizartinib (AC220). **Step 4.** To functionally interpret the two cell – specific causal networks, we searched in SIGNOR and PhosphoSitePlus the direct connections between final network nodes and markers of phenotypes of interest (e.g., BCL2, BAD, etc. were considered markers of apoptosis). Finally, we computed the activity of each marker averaging all the activation state (x_i_) of input nodes multiplied for their regulation sign (sign_i_).

**Figure S6.**
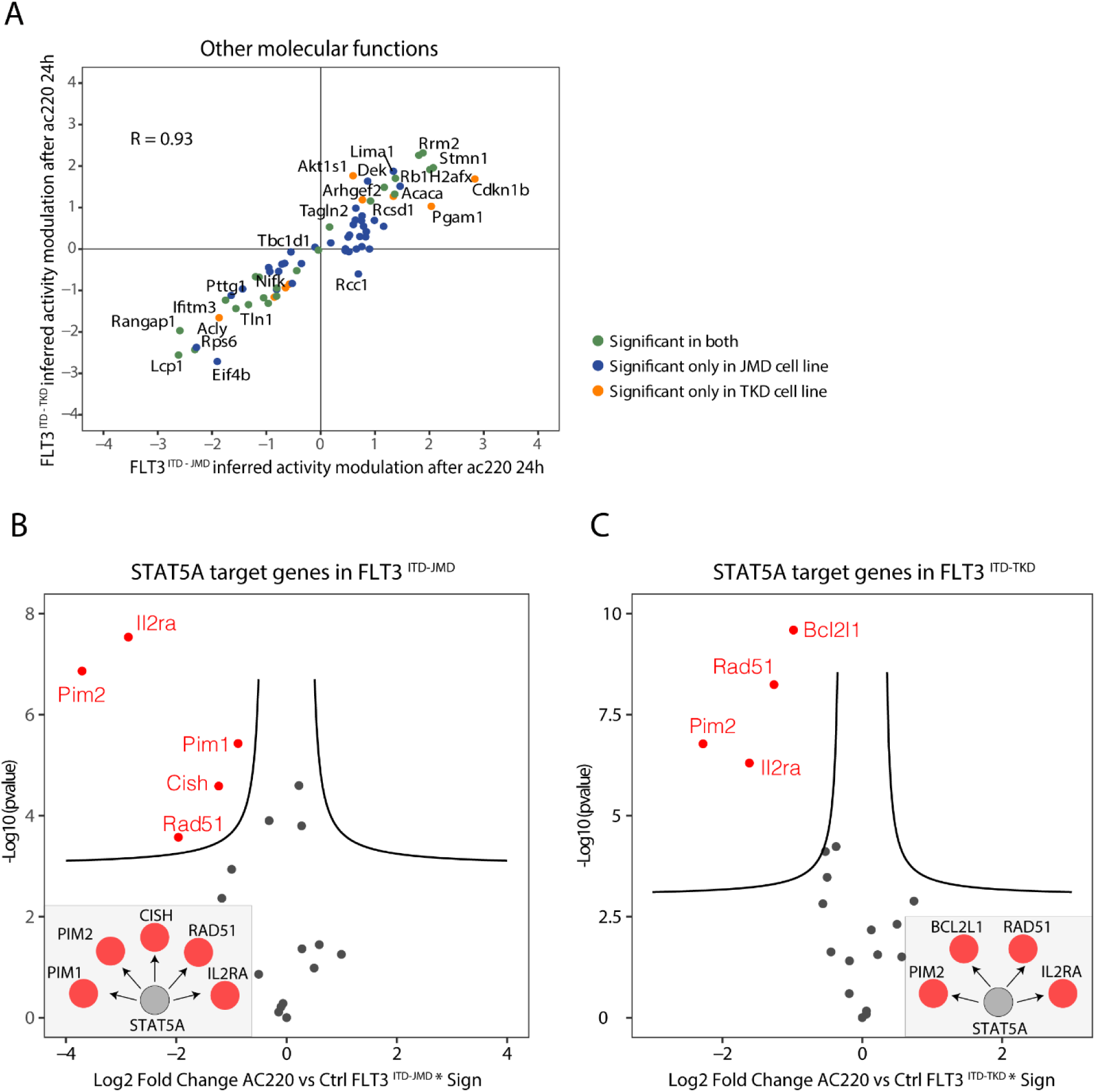
Protein activity prediction from experimental data in quizartinib treated FLT3^ITD^ cells. **(A)** Scatterplot shows the comparison between protein activity predicted from FLT3^ITD-JMD^ (x-axis) and FLT3^ITD-TKD^ (y-axis) datasets for proteins with molecular function different from TF, kinase or phosphatase. Each dot represents a protein, and the color indicates whether the prediction is statistically significant in both cell lines (green) or exclusively in one cell line: FLT3^ITD-JMD^ (blue) or FLT3^ITD-TKD^ (orange). R indicates Pearson correlation. (**B-C**) Volcano plots show the modulation of all STAT5A target genes that were used, by the VIPER tool ^50^, to infer its activity, after quizartinib (AC220) treatment in FLT3^ITD-JMD^ (**B**) and FLT3^ITD-TKD^ cells (**C**). X-axis represents log2 fold change of regulated transcripts multiplied by the sign of regulation (−1 for inhibition, 1 for activation of transcription). Y-axis represents the significance of the log fold change (-log10 pvalue). In each grey panel is shown the regulation of significant transcripts by STAT5A.

**Figure S7.**
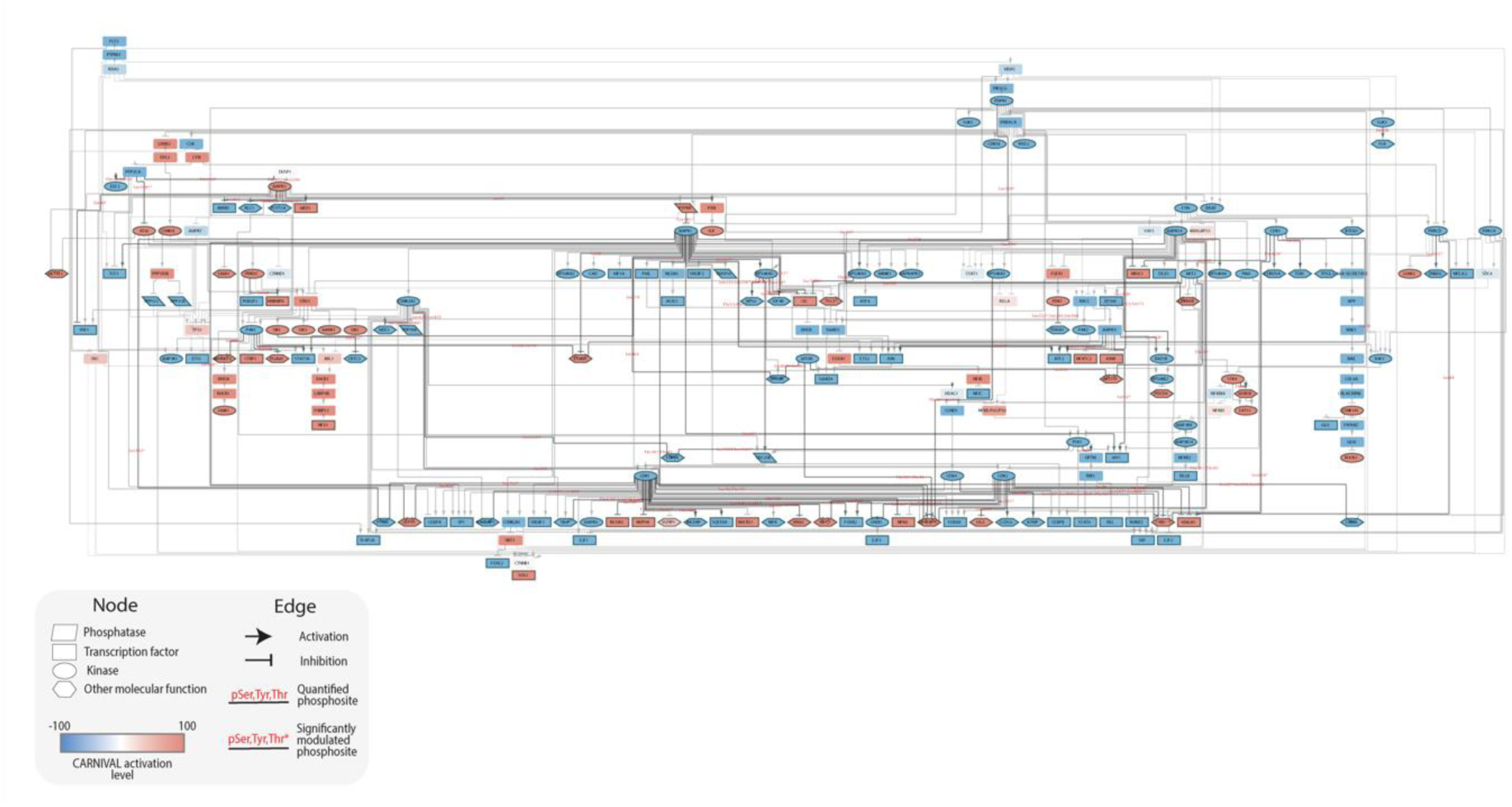
FLT3^ITD-JMD^ specific causal network. Causal network representing the quizartinib-induced signal rewiring in FLT3^ITD-JMD^ cell line. Color of nodes represents activated (red) or inhibited (blue) proteins after the treatment. Shape of nodes reflects molecular function: parallelograms are phosphatases, rectangles are transcription factors, circles are kinases and hexagons are other phosphorylated proteins. Target arrow shape represents activatory (arrow) or inhibitory (T shape) interactions. Black edges represent (de)phosphorylations occurring at phosphosites measured in the experimental data. Additional details are further described in the inset.

**Figure S8.**
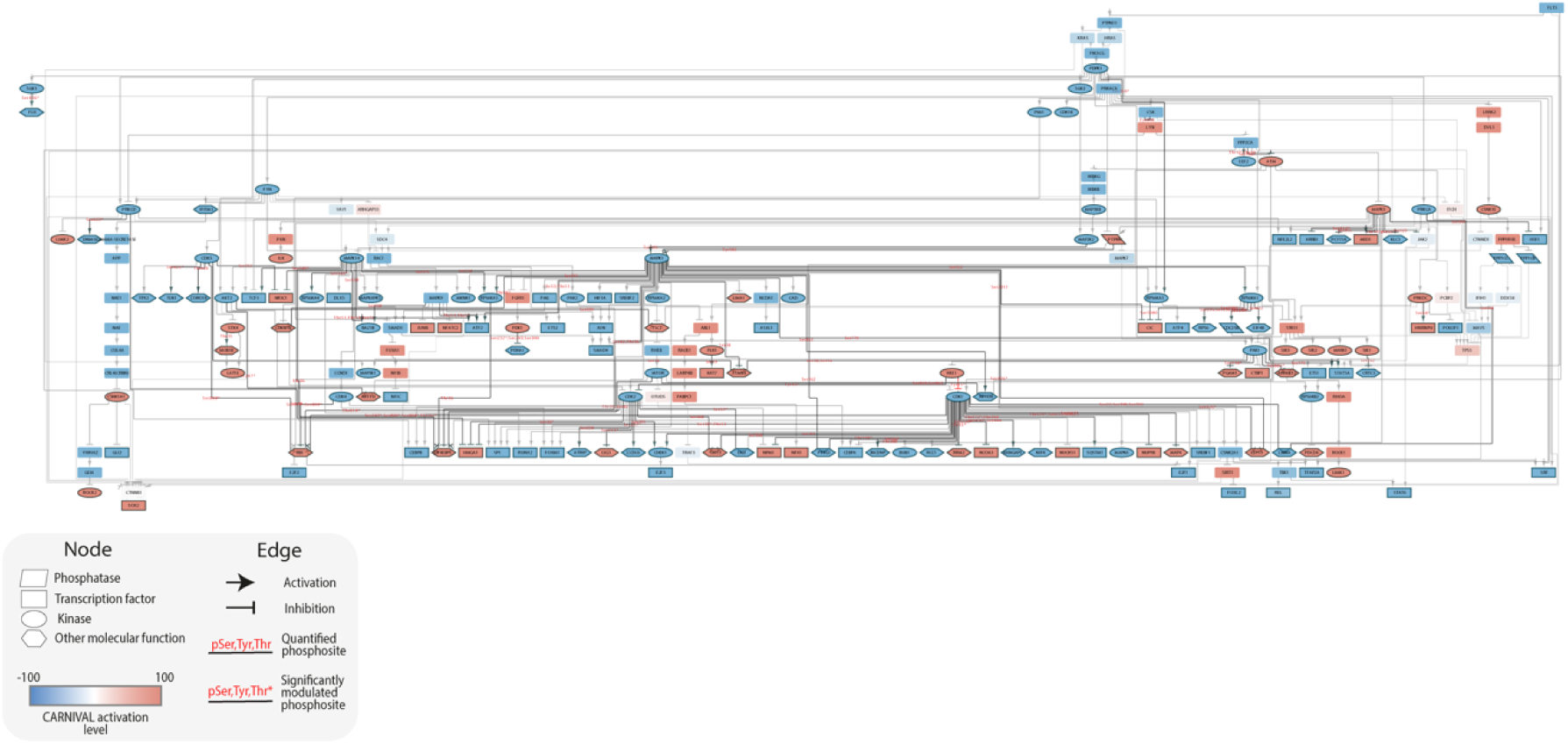
FLT3^ITD-TKD^ specific causal network. Causal network representing the quizartinib-induced signal rewiring in FLT3^ITD-TKD^ cell line. Color of nodes represents activated (red) or inhibited (blue) proteins after the treatment. Shape of nodes reflects molecular function: parallelograms are phosphatases, rectangles are transcription factors, circles are kinases and hexagons are other phosphorylated proteins. Target arrow shape represents activatory (arrow) or inhibitory (T shape) interactions. Black edges represent (de)phosphorylations occurring at phosphosites measured in the experimental data. Additional details are further described in the inset.

**Fig. S9.**
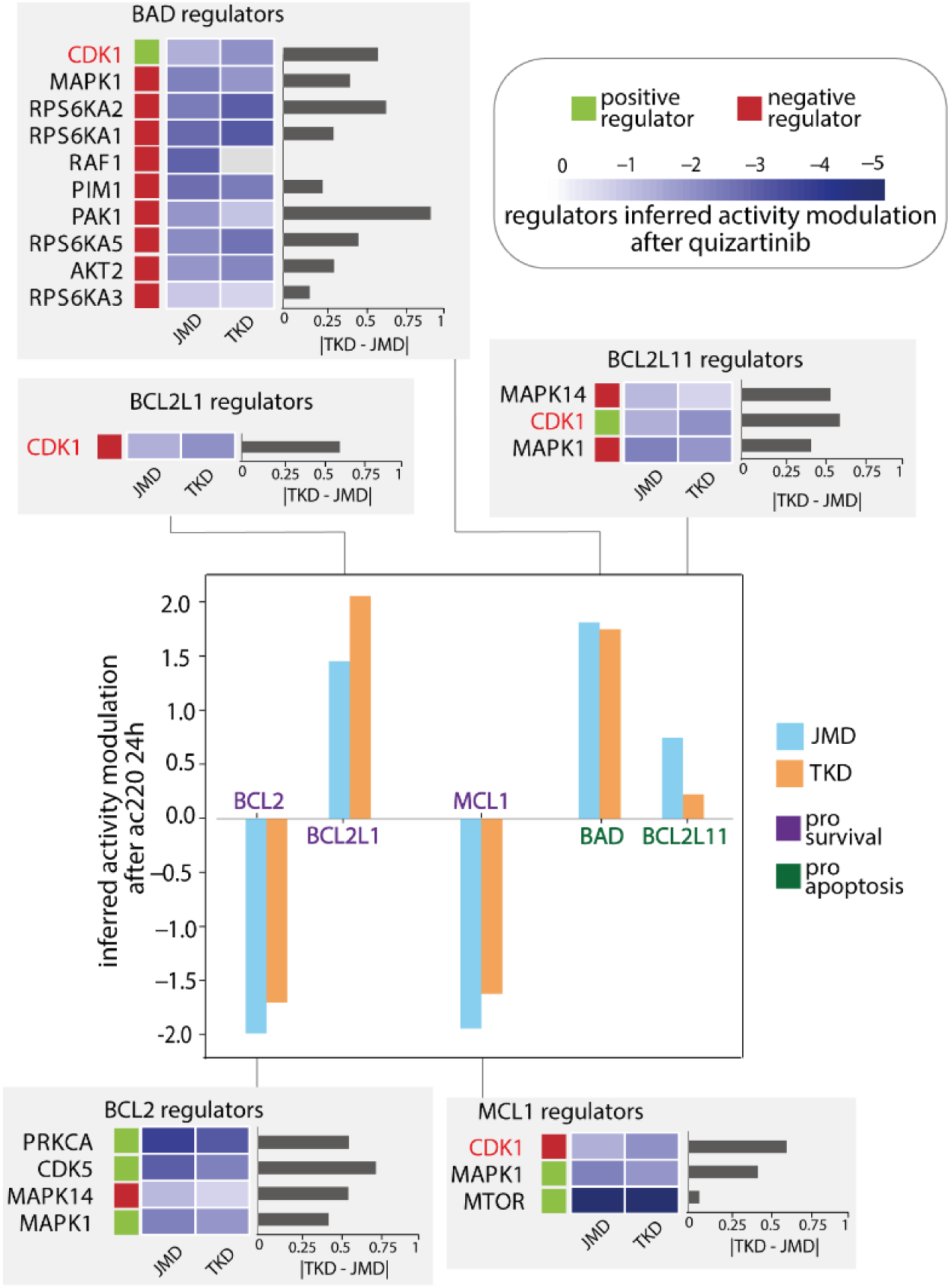
Activity inference of apoptosis markers and activation state of their regulators. Barplot showing the activity score predicted for pro-survival proteins (purple) and pro-apoptotic proteins (green) in FLT3^ITD-JMD^ (blue) and FLT3^ITD-TKD^ (orange) cell lines. In each grey panel, heatmaps show the activation state of upstream regulators of each apoptosis marker. Positive and negative regulators are displayed in green and red, respectively. For each regulator, the barplot on the right reflects the absolute value of the difference in activity between FLT3^ITD-JMD^ and FLT3^ITD-TKD^ models.

**Figure S10.**
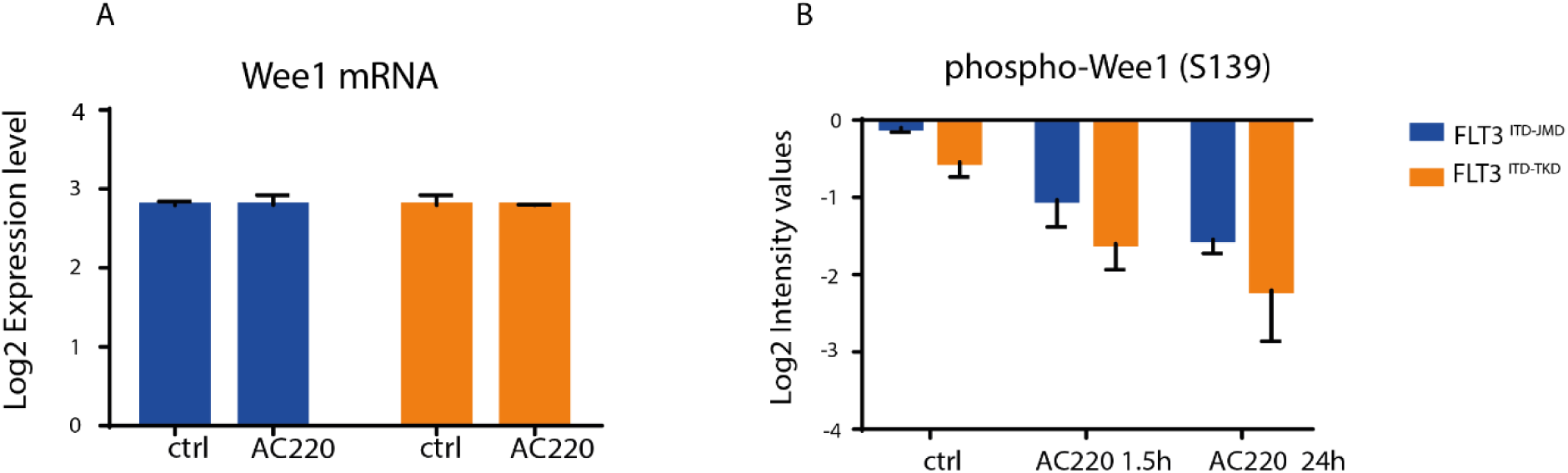
(**A**) Bar plot showing the Log2 expression level of Wee1 mRNA quantified in the transcriptome analysis in FLT3^ITD-JMD^ (blue) and FLT3^ITD-TKD^ (orange) cells after quizartinib (AC220) treatment. (**B**) Bar plot showing the Log2 intensity values of the Wee1 phosphorylation on Serine 139 quantified in the phosphoproteomic analysis after 1.5 hour and 24 hours of quizartinib (AC220) treatment in FLT3^ITD-JMD^ (blue) and FLT3^ITD-TKD^ (orange) cells.

**Figure S11.**
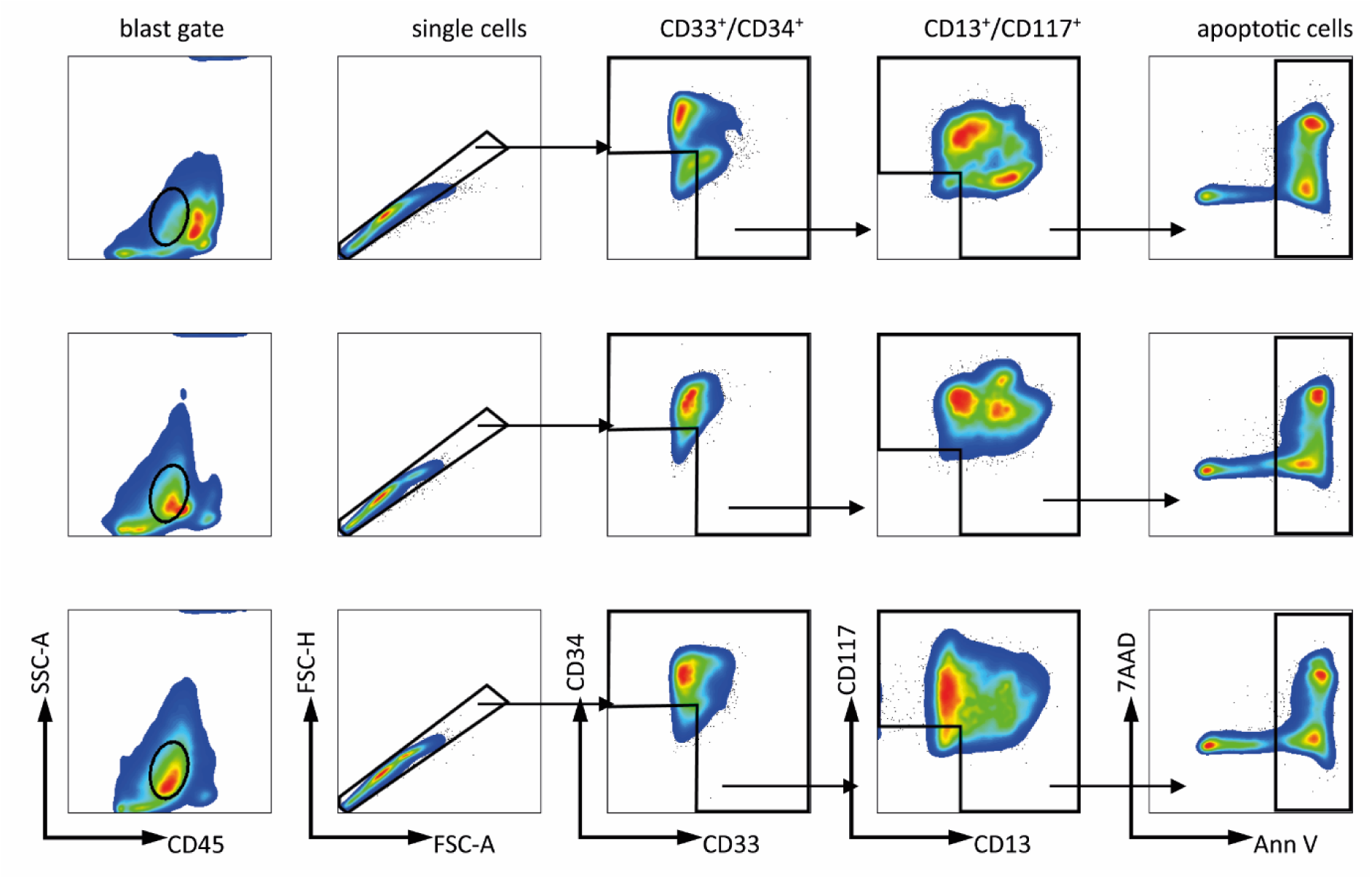
Gating strategy used for the viability assay analysis of primary blasts samples. Blasts were gated based on the CD45/SSC distribution (“blast gate”), for single cells (by FSC-A/FSC-H) and for typical blast markers heterogeneously expressed between patients (CD33, CD34, CD13, CD117). Gates were set according to an unstained/CD45-single stained control. Data was analyzed using FlowJo V10 (Becton-Dickinson, Franklin Lakes, NJ).

**Figure S12.**
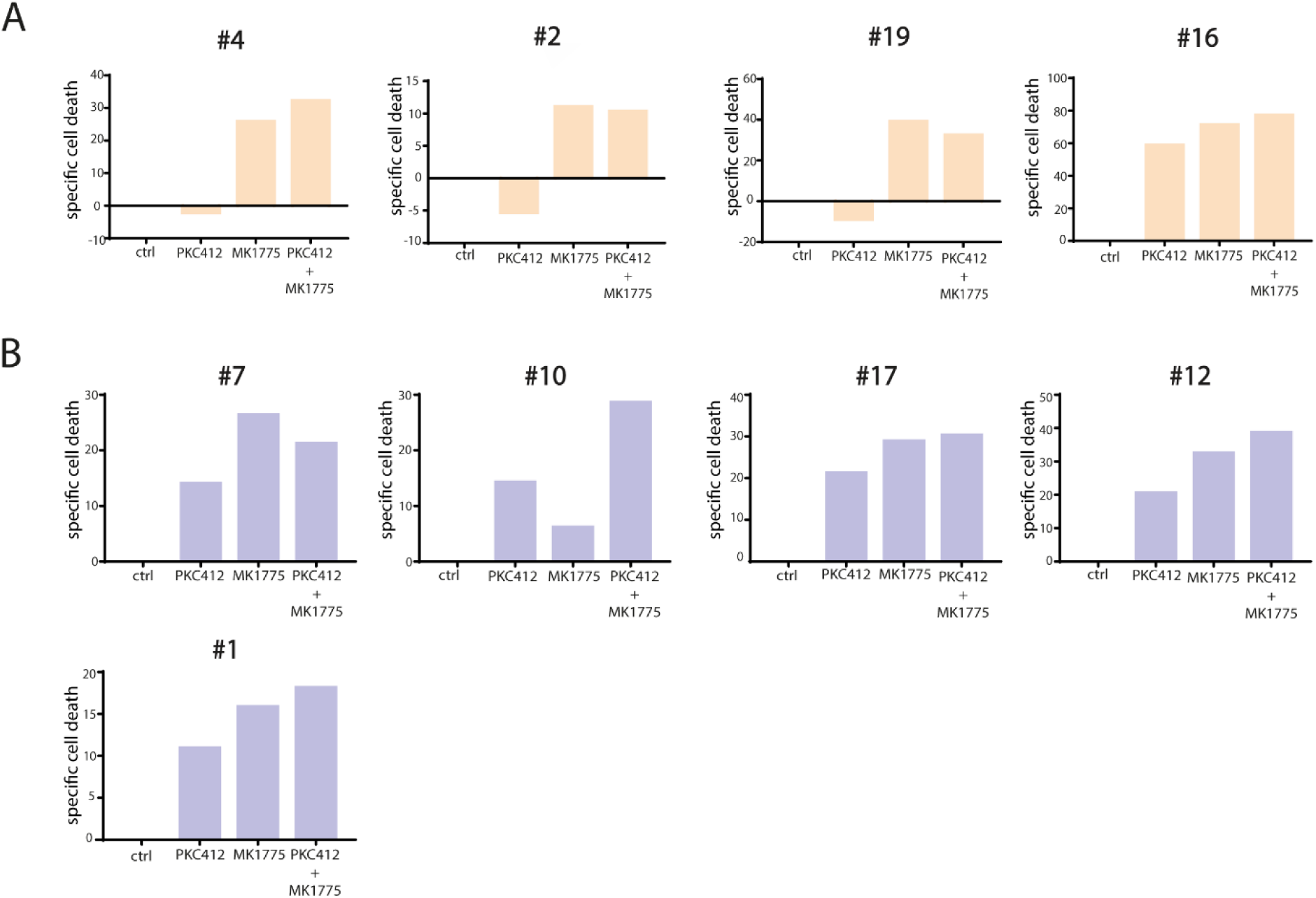
**(A)** Barplots showing blasts treatment specific cell death (*100 * (dead cells after treatment – death cells in control) / viable cells in control*) in each FLT3^ITD-TKD^ patient after 100 nM midostaurin (PRC412), 500 nM Wee1 inhibitor (MK1775) and combination of both. **(B)** Barplots showing blasts treatment specific cell death (*100 * (dead cells after treatment – death cells in control) / viable cells in control*) in each FLT3 ^ITD-JMD+ITD-TKD^ patient after 100 nM midostaurin (PRC412), 500 nM Wee1 inhibitor (MK1775) and combination of both.

